# Towards a Comprehensive Variation Benchmark for Challenging Medically-Relevant Autosomal Genes

**DOI:** 10.1101/2021.06.07.444885

**Authors:** Justin Wagner, Nathan D Olson, Lindsay Harris, Jennifer McDaniel, Haoyu Cheng, Arkarachai Fungtammasan, Yih-Chii Hwang, Richa Gupta, Aaron M Wenger, William J Rowell, Ziad M Khan, Jesse Farek, Yiming Zhu, Aishwarya Pisupati, Medhat Mahmoud, Chunlin Xiao, Byunggil Yoo, Sayed Mohammad Ebrahim Sahraeian, Danny E. Miller, David Jáspez, José M. Lorenzo-Salazar, Adrián Muñoz-Barrera, Luis A. Rubio-Rodríguez, Carlos Flores, Giuseppe Narzisi, Uday Shanker Evani, Wayne E. Clarke, Joyce Lee, Christopher E. Mason, Stephen E. Lincoln, Karen H. Miga, Mark T. W. Ebbert, Alaina Shumate, Heng Li, Chen-Shan Chin, Justin M Zook, Fritz J Sedlazeck

## Abstract

The repetitive nature and complexity of multiple medically important genes make them intractable to accurate analysis, despite the maturity of short-read sequencing, resulting in a gap in clinical applications of genome sequencing. The Genome in a Bottle Consortium has provided benchmark variant sets, but these excluded some medically relevant genes due to their repetitiveness or polymorphic complexity. In this study, we characterize 273 of these 395 challenging autosomal genes that have multiple implications for medical sequencing. This extended, curated benchmark reports over 17,000 SNVs, 3,600 INDELs, and 200 SVs each for GRCh37 and GRCh38 across HG002. We show that false duplications in either GRCh37 or GRCh38 result in reference-specific, missed variants for short- and long-read technologies in medically important genes including *CBS*, *CRYAA*, and *KCNE1*. Our proposed solution improves variant recall in these genes from 8% to 100%. This benchmark will significantly improve the comprehensive characterization of these medically relevant genes and guide new method development.

## Introduction

Authoritative benchmark samples are driving the development of technologies and the discovery of new variants, enabling highly-accurate clinical genome sequencing, and advancing our detection and understanding of the impact of many genomic variations on human disease at scale. With recent improvements in sequencing technologies^1^, assembly algorithms^2–4^, and variant calling methods^5^, genomics offers more insights into challenging genes associated with human diseases across a higher number of patients^6^. Still, challenges remain for medically-relevant genes that are often repetitive or highly polymorphic^7,8^. In fact, a recent study found 13.8 % (17,561) of pathogenic variants identified by a high-throughput clinical laboratory were challenging to detect with short-read sequencing^9^. These included challenging variants such as variants 15 bp to 49 bp in size, small copy number variations (CNVs), complex variants, as well as variants in low-complexity or segmentally duplicated regions.

The Genome in a Bottle (GIAB) consortium develops benchmarks and standards to advance accurate human genomic research and clinical applications of sequencing. GIAB provides highly-curated benchmark sets for single-nucleotide variant (SNV)^10^, small insertion and deletion (INDEL)^10^, and structural variant (SV) calling^11^. Here, we define SNVs as single base substitutions, while INDELs are defined as insertions and deletions smaller than 50 bp, in contrast to insertions and deletions larger than 50 bp, which we refer to as SVs. Furthermore, GIAB and the FDA host periodic precisionFDA challenges providing an important snapshot and recommendations for small variant calling enabling the high precision and sensitivity required for clinical research, with a recent challenge demonstrating the importance of including more difficult genomic regions^12^. Recently, GIAB focused primarily on a read mapping based genome-wide approach integrating short-, linked-, and long-read sequencing to characterize up to 92% and 86% of the autosomal bases for small variants and SVs, respectively^11,13^. GIAB also released a targeted assembly-based benchmark for the MHC region, a highly diverse and repetitive region of the human genome that includes the HLA genes^14^. Still, multiple regions of the genome are not fully resolved in existing benchmarks due to repetitive sequence, segmental duplications, and complex variants (i.e., multiple nearby SNVs, INDELs, and/or SVs)^15^.

Interestingly, many clinically-relevant genes are in the remaining hard-to-assess regions. The clinical tests for these genes often require locus-specific targeted designs and/or employ multiple technologies, and are only applied when suspicion for a specific disorder is high. Mandelker *et al*. categorized genes based on their repetitive content and identified 193 genes that cannot be fully characterized by short-read sequencing^7^. This gene set was constructed by identifying genes with low mapping quality in the clinical databases OMIM, HGMD, and ClinVar. Subsequently, Wenger *et al*. showed that, while short reads could not accurately map the full length of these genes, highly-accurate long-reads could fully map 152 (78.76%) of them^1^. The latest v4.2.1 GIAB small variant benchmark regions included at least 90% of the gene body for 110 out of the 159 difficult genes on autosomes.^13^ In contrast, the previous v3.3.2 GIAB small variant benchmark regions included at least 90% of the gene body for only 19 out of 159 difficult genes.^10^ Although v4.2.1 includes substantially more difficult genes, variant calls in the remaining most difficult genes still need to be assessed, and challenges remain with typical mapping-based approaches in some genes even when using highly accurate long reads.

To support ongoing advancements in clinical genome sequencing and bioinformatics, we present a more comprehensive benchmark of challenging, medically-relevant genes (CMRG) focusing on HG002, who has a broad consent from the Personal Genome Project for open genomic data and commercial redistribution ^16^ (**Figure 1**). With the advent of highly-accurate long reads, new approaches for haplotype-resolved (diploid) assembly have advanced rapidly^2,3^. Here, we focus on generating a benchmark for as many of these genes as possible, the first benchmark using a whole genome diploid assembly. We curated a set of 273 medically-relevant genes with <=90% bases included in previous GIAB benchmarks but fully covered by both haplotypes of a trio-based hifiasm assembly. The assembly included all phased small variants and SVs across these genes. Then, we delineated regions where we can provide reliable small variant and SV benchmarks, developing a prototype process for future whole genome assembly-based benchmarks. Overall, this represents an important step in the effort to drive clinical genetics forward by providing an accurate benchmark for medically relevant genes that have so far escaped complete characterization.

**Figure 1:**
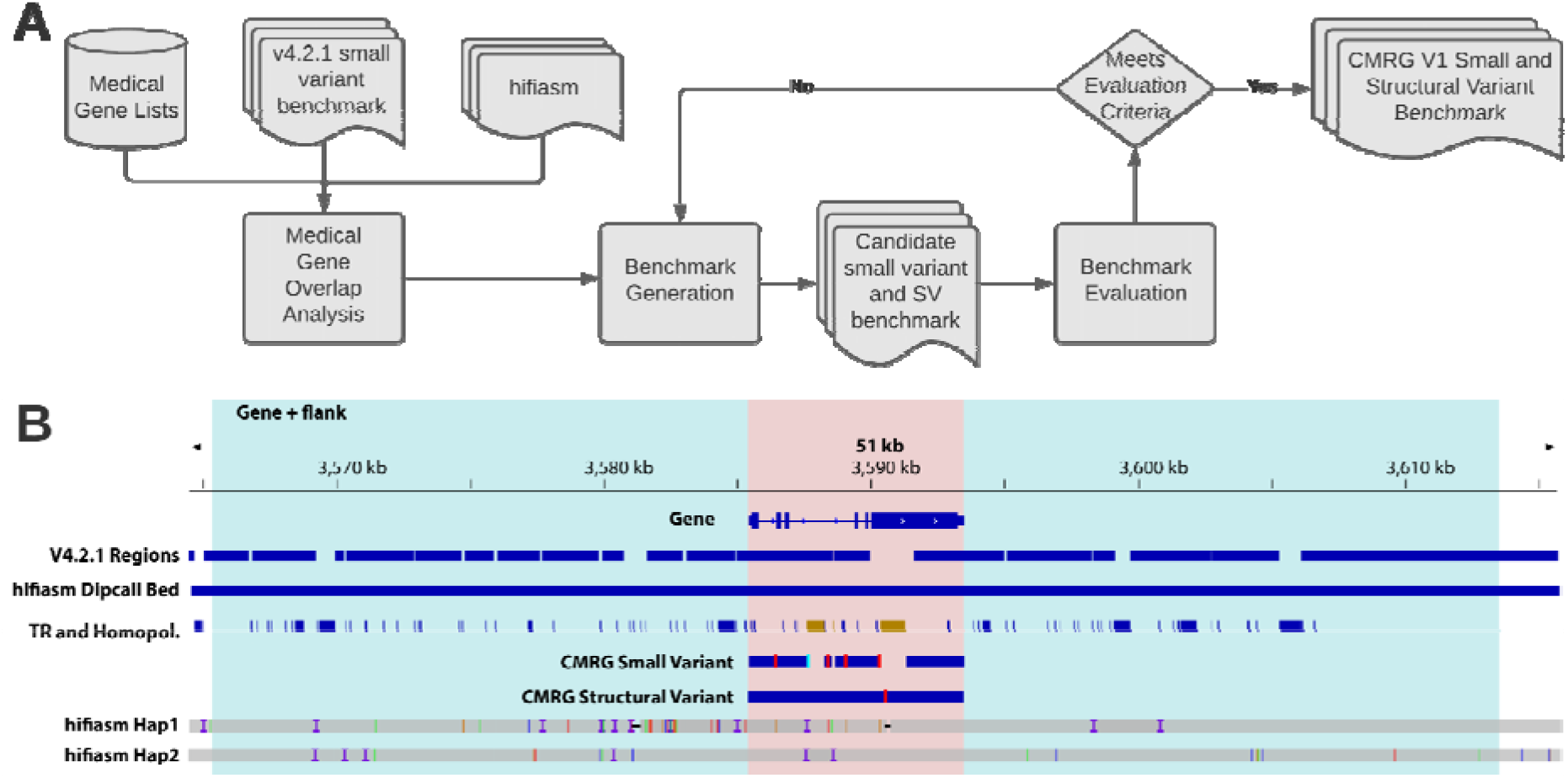
GIAB developed a process to create new phased small variant and structural variant benchmarks for 273 challenging, medically relevant genes. (A) We developed a list of 4,701 autosomal potentially medically relevant genes. We generated a new benchmark for 273 of the 4,701 genes that were completely resolved by our hifiasm diploid assembly and <=90% included in the v4.2.1 GIAB small variant benchmark for HG002 (V4.2.1 Regions). (B) We required that the entire gene region (pink) and 20 kb flanking sequence on each side (blue) were completely resolved by both haplotypes in the assembly (hifiasm Hap1 and hifiasm Hap2), indicated with the hifiasm Dipcall Bed track. In addition, we required that any segmental duplications overlapping the gene were completely resolved. From the small variant benchmark regions (CMRG Small Variant blue bars), we excluded SVs and any tandem repeats or homopolymers overlapping SVs (right TR and Homopol. region in brown). The left TR and Homopol. region in brown is excluded from the small variant benchmark regions because the larger tandem repeat contains an imperfect homopolymer longer than 20 bp, which we exclude because long homopolymers have a higher error rate in the assembly. All regions of this gene were included in the SV benchmark regions (CMRG Structural Variant blue bar). The vertical red lines in CMRG Small Variant and CMRG Structural Variant indicate locations of benchmark small variants and SVs, respectively. Finally, we evaluated the small variant and structural variant benchmarks with manual curation and long range PCR, and also ensured they accurately identify false positives and false negatives after excluding errors found during curation.

## Results

### Identification of challenging, medically relevant genes

To prioritize genome regions for the new expanded benchmark, we identified several lists of potentially medically-relevant genes. (1) 4,773 potentially medically-relevant genes from the databases OMIM, HGMD, and ClinVar previously compiled in 2012, which includes both commonly tested and rarely tested genes (Supplementary Table 13 in ^7^). (2) The COSMIC gene census contains 723 gene symbols found in tumors (https://cancer.sanger.ac.uk/census)^17^. (3) We developed a focused list of “High Priority Clinical Genes” that are commonly tested for clinical inherited diseases (**Supplementary File 1**). There are 5,175 gene symbols in the union of these sets, of which 5,027 have unique coordinates on the primary assembly of GRCh38 and valid ENSEMBL annotations, and 4,697 are autosomal. 70% of these genes are specific to the list from OMIM, HGMD, and ClinVar, which includes genes associated with disease in a small number of studies and are currently tested more frequently in research studies than in high-throughput clinical laboratories (**Figure 1(A)**).

To identify genes for which a new benchmark was most needed, we next examined the fraction of the 4,697 autosomal medically-relevant gene bodies included in the latest GIAB HG002 v4.2.1 small variant benchmark. We excluded the sex chromosomes, X and Y, because v4.2.1 did not include these haploid chromosomes in males like HG002. The median fraction of each autosomal gene included in v4.2.1 is 97.8%, with 544 (11.6%) genes completely included in v4.2.1, 3,746 included > 95%, and 4,343 (92.5%) included > 90% (**Supplementary File 2**). Genes were not fully included in v4.2.1 for a variety of reasons, including: (1) putative SVs, specifically regions containing GIAB’s HG002 v0.6 Tier 1 or Tier 2 SVs^11^, some of which are true SVs and some of which do not contain SVs or only partly contain SVs, (2) regions difficult to characterize with mapping-based methods, including complex variants in segmental duplications and long tandem repeats that were not properly resolved with mapping of Illumina or PacBio HiFi reads, (3) large duplications in HG002 not in GRCh38, in which small variants cannot be represented in standardized ways, and (4) variants (plus 50 bp flanking regions) that were not considered high confidence by the GIAB small variant integration pipeline because differences between methods could not be resolved, or all methods had evidence of bias. We identified 361 and 354 autosomal genes that are included 90% or less by the HG002 v4.2.1 benchmark regions on GRCh37 or GRCh38, respectively, totaling 395 unique genes. We focused this work on genes previously 90% or less included in order to facilitate our manual curation of the genes, thoroughly inspecting variant calls from the diploid assembly and the accuracy of the assembly for these challenging medically relevant genes (CMRG). Diploid assembly enables a phased small and structural variant benchmark for 273 challenging medically relevant genes

Many of the 395 medically relevant genes were not covered well by the v4.2.1 small variant benchmark due to SVs, complex variants, and segmental duplications (**Figure 2**). Thus, here we resolve many of these regions using a diploid assembly of HG002 constructed by hifiasm^2^. This approach constructs a graph using HiFi reads from HG002 and separates the parental haplotypes based on unique kmers identified in short reads of the mother and father. Hifiasm can resolve both haplotypes with high base-level quality (QV>50), including many segmental duplications, and produces variant calls and genotypes that are highly concordant with the v4.2.1 small variant benchmark, with both recall and precision >99.7% for SNVs and >97% for INDELs in regions covered by the assembly (**Supplementary File 3**).

**Figure 2.**
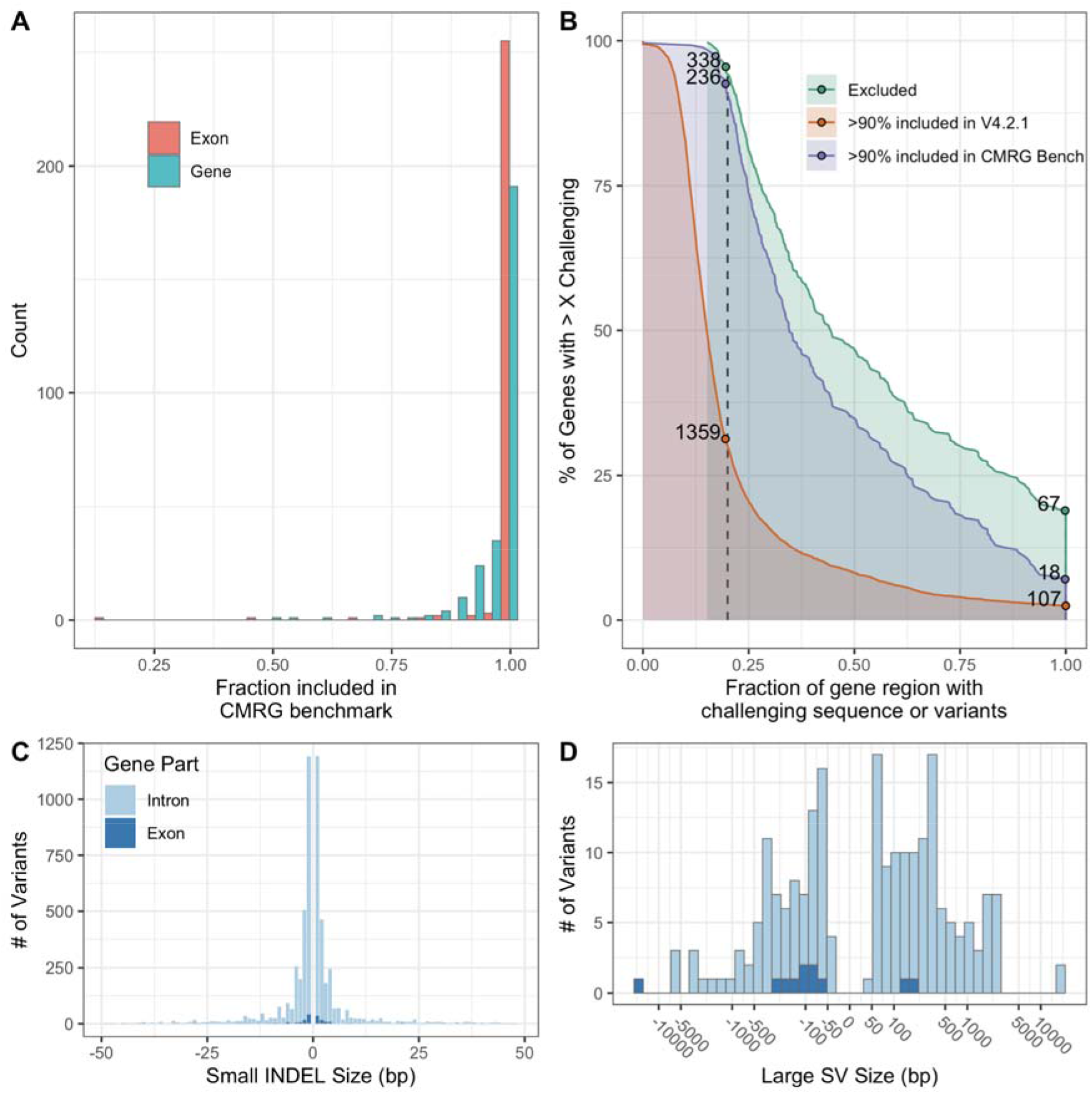
The new CMRG benchmark contains more challenging variants and regions than previous benchmarks. (A) Fraction of each gene region (blue) and exonic regions (red) included in the new CMRG small variant or SV benchmark regions. (B) Comparison of fraction of challenging sequences and variants for genes included in the new CMRG benchmark vs. the previous v4.2.1 HG002 benchmark vs. genes excluded from both benchmarks. 99% of CMRG benchmark genes have at least 15% of the gene region with challenging sequences or variants. The catalog of repetitive challenging sequences comes from GIAB and the Global Alliance for Genomics and Health (see text). Challenging variants for HG002 are defined as complex variants (i.e., more than one variant within 10 bp) as well as putative SVs and putative duplications excluded from the HG002 v4.2.1 benchmark regions. C) Size distribution of INDELs in the small variant benchmark, which includes some larger INDELs in introns (light blue) and exons (dark blue). D) Size distribution of large insertions and deletions in the SV benchmark in introns (light blue) and exons (dark blue).

We generated a focused benchmark (see **Methods**) for 273 of the 395 genes that were fully resolved by this assembly. To be included in the CMRG benchmark, the entire gene including 20 kb flanking sequence on each side and any overlapping segmental duplications needed to have exactly one fully aligned contig from each haplotype with no breaks on GRCh37 and GRCh38 (**Supplementary File 2**). We required the alignment to include flanking sequences greater than the size of the longest reads used by hifiasm, and that the alignments completely resolve any overlapping segmental duplications to minimize ambiguity or errors in the assembly-assembly alignment. These 273 genes are substantially more challenging than genes previously covered by GIAB’s v4.2.1 benchmark; for example, for 99% of the new genes, at least 15% of the gene region is either challenging to sequence or contains challenging variants in HG002 (**Figure 2(B)**). Here, we use the definition of challenging sequences from GIAB and the Global Alliance for Genomics and Health:^12^ specifically, the union of all tandem repeats, all homopolymers >6 bp, all imperfect homopolymers >10 bp, all difficult to map regions, all segmental duplications, GC <25% or >65%, GC-rich promoters that are difficult to sequence,^18^ regions near gaps in the reference, the MHC, and the immunoglobulin V(D)J region. Furthermore, when comparing variants in regions of the CMRG gene bodies included by the v4.2.1 benchmark, the CMRG benchmark, or both benchmarks, 11% of the CMRG benchmark INDELs are >15 bp (**Figure 2(C)**, vs. 3.5% in v4.2.1. The CMRG INDELs >15 bp are also substantially more challenging than the v4.2.1 INDELs >15 bp. This is shown by a 17.2% decrease in recall of HiFi-Deep Variant from 99.5% (v4.2.1) to the new CMRG (84.9%). In addition, the precision has decreased by 3.9% for HiFi-DeepVariant from 99.9% (v4.2.1) to 94.2% (CMRG) (**Supplementary File 4**).

An important step in benchmark formation is excluding regions where the benchmark is unreliable or has variants that cannot be compared robustly with current benchmarking tools (e.g., regions with complex SVs that can have different representations). No benchmarking tools currently exist to jointly benchmark small variants and SVs. Therefore, we created separate CMRG benchmark bed files for small and structural variants, which both rely on the same benchmark variant calls from hifiasm. The CMRG benchmark extends beyond the v4.2.1 benchmark across the 273 challenging gene regions, adding many new phased SNVs, INDELs, and large insertions and deletions at least 50 bp in length overlapping these genes (**Table 1**). With this approach, we resolved these challenging but relevant genes completely across GRCh37 and GRCh38.

**Table 1:** Number of bases and variants in different HG002 GIAB benchmarks sets included in the 273 genes in the CMRG benchmark. We denote the percent of base pairs or variants across exons in the brackets. Difficult context defined as union of all tandem repeats, all homopolymers >6 bp, all imperfect homopolymers >10 bp, all difficult to map regions, all segmental duplications, GC <25% or >65%, “Bad Promoters”, and “OtherDifficultregions”. Challenging variants for HG002 are defined as complex variants (i.e., more than one variant within 10 bp) as well as putative SVs and putative duplications excluded from the HG002 v4.2.1 benchmark regions. Number of bases and variants are provided for benchmarks on GRCh38, except for v0.6 where only a GRCh37 benchmark is available.

| Benchmark Set | bp in CMRG Benchmark Genes | bp in Difficult Context | # of Variants |
| --- | --- | --- | --- |
| CMRG Small Var. | 11 719 200 (11.5 %) | 4 500 129 (13.4 %) | 27 178 ( 5.2 %) |
| v4.2.1 Small Var. | 9 763 722 (12.0 %) | 2 637 132 (16.3 %) | 16 804 ( 6.3 %) |
| CMRG SV | 12 020 518 (11.4 %) | 4 792 096 (13.0 %) | 217 ( 5.1 %) |
| v0.6 SV | 10 569 811 ( 9.6 %) | 3 215 766 (13.1 %) | 170 ( 4.7 %) |

### Resolving Challenging Medically Relevant Genes

Beyond previous GIAB benchmarks, this new CMRG benchmark improves upon 273 important, more challenging genes. These include (1) genes that are duplicated in the reference but not in HG002, as described above, (2) highly homologous genes such as *SMN1* and *SMN2* or *NCF1*, *NCF1B*, and *NCF1C*, and (3) genes with SVs and complex variants like *RHCE*.

The important gene *SMN1* resides within a large segmental duplication on chromosome 5 containing both *SMN1* and *SMN2*. Biallelic pathogenic variants in *SMN1* result in spinal muscular atrophy (SMA), a progressive disorder characterized by muscle weakness and atrophy due to loss of neuronal cells in the spinal cord^19^. While the 28 kb sequences of *SMN1* and *SMN2* generally differ by only 5 intronic and 3 exonic nucleotides^20^, the identification and characterization of pathogenic variants in *SMN1* and copy-number state of *SMN2* is important in guiding newly developed therapies and counseling families regarding recurrence risk of this disease. Although *SMN2* has common copy number polymorphisms, HG002 appears to contain one copy each of *SMN1* and *SMN2* on each haplotype based on the presence of two haplotypes for each gene in ONT and 10x Genomics data. However, the genes are surrounded by complex repeats and are thus not both fully resolved by our assembly (**Figure 3(B)**). The maternal assembly has a single contig passing through the SMA region but misses *SMN2* and some of the surrounding repeats (dot plot in **Supplementary Figure 1**). The paternal assembly contains both *SMN1* and *SMN2* but the assembly is broken into three contigs in the SMA region (dot plot in **Supplementary Figure 1**). Upon curation of the data from PacBio HiFi, ultralong ONT, and 10x Genomics in **Figure 3(A)**, the variants called from the assembly of *SMN1* were supported by ONT and 10x Genomics across the full gene and by PacBio HiFi across the part of the gene covered by reads. Because we manually confirmed the assembly accuracy in this important gene, we included *SMN1* in our benchmark even though our general heuristics excluded it because the assemblies did not cover the segmental duplications within the entire SMA region. We excluded *SMN2* because only one haplotype was resolved by hifiasm v0.11. Another example is *NCF1*, which is associated with 20% of cases of chronic granulomatous disease (CGD), a primary immunodeficiency^21^. The gene lies within a large segmental duplication, which may make molecular diagnosis of some cases of CGD challenging. The new benchmark covers the first two exons that were missing from the v4.2.1 benchmark (**Supplementary Figure 2**).

**Figure 3:**
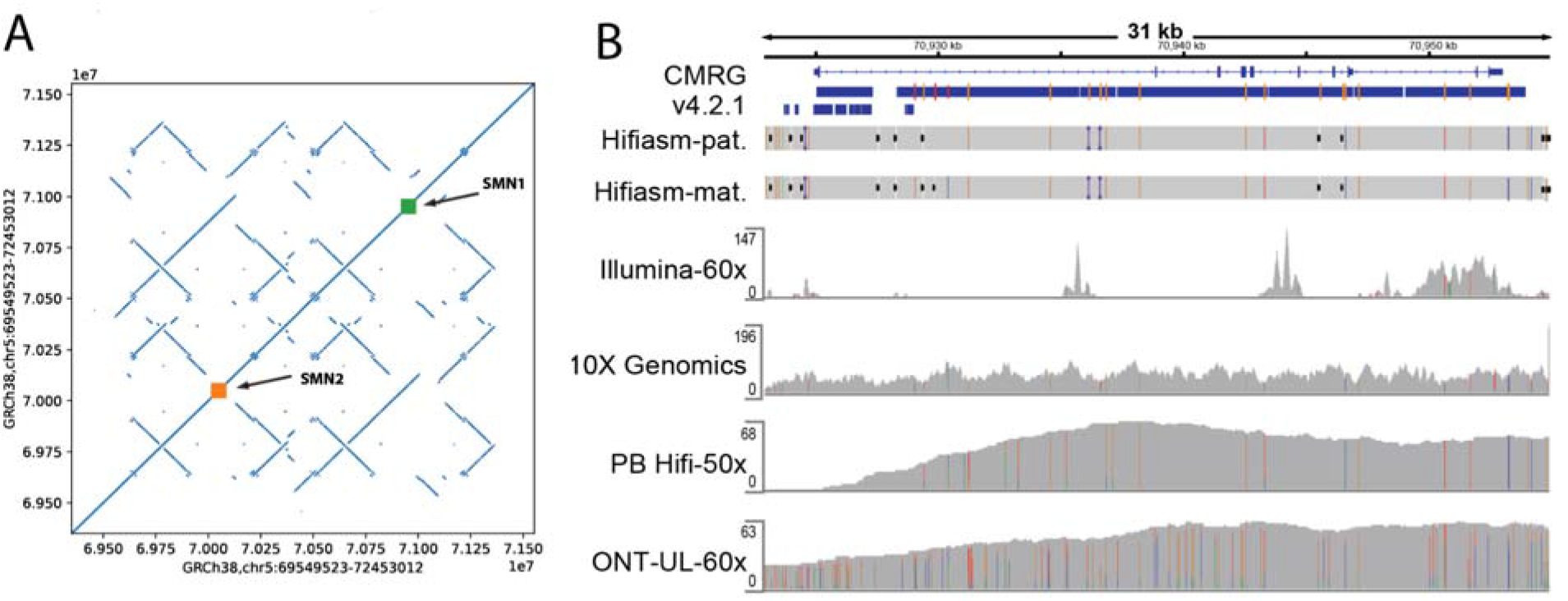
The new benchmark covers the gene *SMN1*, which was previously excluded due to mapping challenges for all technologies in the highly identical segmental duplication. (A) Dotplot of GRCh38 against GRCh38 in the SMA region, showing a complex set of inverted repeats that make it challenging to assemble. (B) IGV view showing that only a small portion of *SMN1* was included in v4.2.1, and that all technologies have challenges mapping in the region, but 10x Genomics and ultralong ONT reads support the variants called in the new CMRG benchmark. For the CMRG and v4.2.1 benchmarks, thick blue bars indicate regions included by each benchmark and orange and light blue lines indicate positions of homozygous and heterozygous benchmark variants, respectively. CMRG variants were called from the trio-based hifiasm assembly of paternal and maternal haplotypes (Hifiasm-pat and Hifiasm-mat, respectively). Coverage tracks are show for 60x PCR-free Illumina 2×150 bp reads (Illumina-60x), 10x Genomics linked reads (10X Genomics), 50x PacBio HiFi 15 kbp and 20 kbp reads (PB Hifi-50x), and 60x Oxford Nanopore ultralong reads (ONT-UL-60x).

HG002 has an approximately 4.5 kb region with an SV and many homozygous small variants surrounding exon 2 of *RHCE*, a gene that is part of the Rh blood group antigens. *RHCE* and *RHD* are part of an inverted duplication on chromosome 1. This complex set of variants results from a 4.5 kb region of *RHCE* in both HG002 haplotypes that have a sequence very similar to a 4.5 kb region of *RHD* in GRCh37 and GRCh38, which may be related to past gene conversion events^22^. While long reads align correctly to both genes, short reads and linked reads that should align to this region in *RHCE* incorrectly align to *RHD*, leading to low coverage in *RHCE* (**Supplementary Figure 3**) and high coverage in *RHD* (in the region chr1:25,283,400-25,287,900 on GRCh38 in **Supplementary Figure 4**), as well as some discordantly paired reads between the genes. Our benchmark provides a way to measure accuracy of variant calls in gene conversion-like events, though performance may differ depending on the characteristics of the gene conversion (e.g., for gene conversions between the highly identical *SMN1* and *SMN2* genes). A similar interesting gene from a technical perspective with common short read-based false negatives is *SIGLEC16*. *SIGLEC16* has evolutionarily gone through several gene conversion events with nearby *SIGLEC11^23^* and one copy of *SIGLEC16* in HG002 has a dense series of variants that causes a drop in short-read coverage.

### CMRG benchmark for challenging structural variants

This curated CMRG benchmark includes new, more challenging classes of SVs not included in the previous GIAB v0.6 benchmark: (1) It includes a sequence-resolved, large 16,946 bp insertion in a VNTR (variant number tandem repeat) in an intron of the gene *GPI*, which is challenging to call with mapping-based methods, even with long reads (**Supplementary Figure 5(a))**. Although VNTRs have been difficult to study, recent evidence points to association of VNTRs with methylation and gene expression^24^. (2) It includes SVs in segmental duplications such as a homozygous 2.3 kb intronic VNTR expansion in *PKD1*, a 5.9 kb homozygous LINE:L1HS deletion in *SMG1*, and two homozygous insertions (588 bp and 1,205 bp) in the gene *GTF2IRD2* on GRCh38 (**Supplementary Figure 5(b))**. The two insertions in *GTF2IRD2* are often missed by mapping-based variant callers because even long reads mismap to the other copy of the segmental duplication, which contains sequences similar to the inserted sequences. GRCh38 corrected a tiling path issue in GRCh37 that mixed haplotypes and resulted in a gap in the region. Interestingly, this resulted in a different representation of *GTF2IRD2*, such that instead of the 588 bp and 1,205 bp insertions, HG002 has a homozygous 195 bp deletion relative to GRCh37, but many short and long reads in this region still mismap to the other copy of the segmental duplication. (3) It includes compound heterozygous insertions (i.e., each haplotype has a different large insertion size in a tandem repeat). (4) Other complex SVs like those in *FLG* and *DSPP* are included in the benchmark VCF, but excluded by the benchmark regions because of a lack of tools to do the comparison robustly. Nevertheless, these SVs that cannot be benchmarked with automated tools could be compared manually or with future benchmarking tools. In addition, the SV sizes and estimated insertion and deletion sequences in the new SV benchmark are more accurate than v0.6 because v0.6 did not include accurate HiFi reads or trio-based partitioning of haplotypes for assembly. The small variant benchmark VCF also contains phased SNVs and INDELs near the benchmark SVs, though these are excluded from the small variant benchmark regions. These large SVs, complex SVs, and SVs inside segmental duplications enable benchmarking of more challenging SVs than previously possible.

### Identifying and resolving false duplications of important genes in the reference

The CMRG benchmark identified variant calling errors due to false duplications in GRCh37 or GRCh38 in several medically-relevant genes. Previous work described true highly homologous genes inside segmental duplications in GRCh37 and GRCh38 that give rise to read mapping issues^7,8^; our new CMRG benchmark, however, identifies that several of these highly homologous genes are in fact false duplications in the reference. For example, PacBio HiFi and Illumina short-read coverage is low and missing one or both haplotypes for *CBS, CRYAA,* and *KCNE1* on GRCh38 because reads incorrectly align to distant incorrect copies of these genes - *CBSL*, *CRYAA2*, and *KCNE1B*, respectively (**Figure 4, Supplementary Figure 6, and Supplementary Figure 7).** Clarification of these regions is important, such as for *CBS*, whose deficiency is associated with homocystinuria, a disorder associated with thromboembolic events, skeletal abnormalities, and intellectual disability. Most cases of homocystinuria are detected by newborn screening and subsequent molecular evaluation can help confirm the diagnosis and provide important recurrence risk information for families of affected individuals. *H19*, a non-coding gene on chromosome 11 that is frequently evaluated in cases of Beckwith-Wiedemann syndrome^25^, is similarly affected by a false duplication on GRCh38. The extra copies of *CBS, U2AF1, CRYAA,* and *KCNE1* in GRCh38 do not occur in HG002, and the Genome Reference Consortium and Telomere to Telomere Consortium recently determined that several regions on the p arm of chromosome 21 as well as several other regions in GRCh38 were incorrectly duplicated^26,27^. In support of this, the gnomAD v2 database has normal coverage and variants called in these genes for GRCh37, but gnomAD v3 has very low coverage and few variants for GRCh38. A companion manuscript from the Telomere-to-Telomere Consortium demonstrates that the new T2T-CHM13 reference corrects these and additional false duplications affecting 1.2 Mbp and 74 genes.^27^

**Figure 4:**
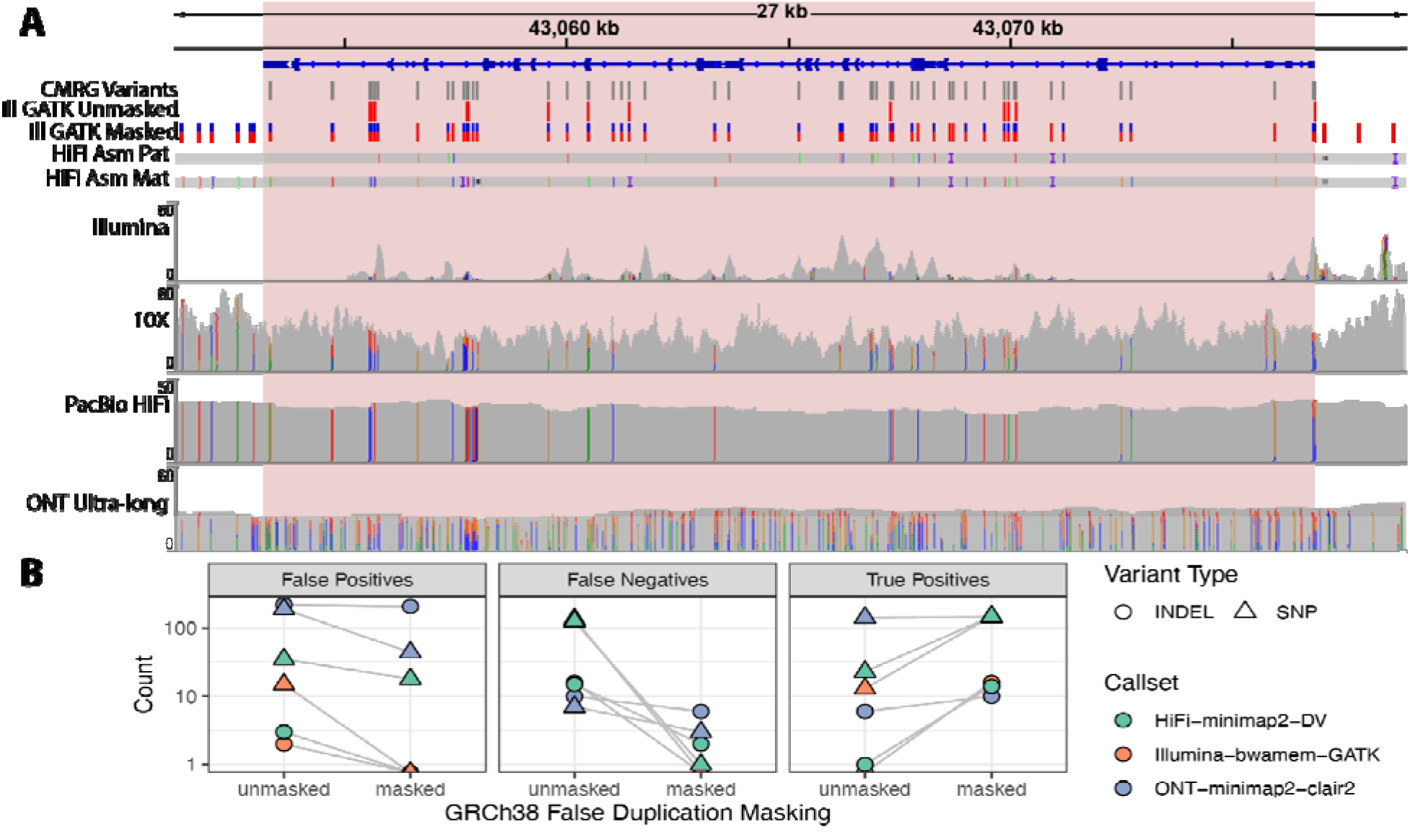
(A) The benchmark resolves the gene *CBS*, which has a highly homologous gene *CBSL* due to a false duplication in GRCh38 that is not in HG002 or GRCh37. The duplication in GRCh38 causes Illumina and PacBio HiFi reads from one haplotype to mismap to *CBSL* instead of *CBS*. The ultralong ONT reads, 10x Genomics linked reads, and assembled PacBio HiFi contigs map properly to this region for both haplotypes because they contain sufficient flanking sequence. When the falsely duplicated sequence is masked using our new version of GRCh38, variant calls from a standard Illumina-GATK pipeline (ILMN-GATK w/ Mask VCF) are completely concordant with the new benchmark. Pink shaded box indicates CMRG benchmark regions, only variants within the benchmark regions are included in the benchmark. (B) Comparison of variant accuracy for GRCh38 before and after masking false duplications on chromosome 21. The new benchmark demonstrates decreases in false negative and false positive errors for 3 callsets in the falsely duplicated genes *CBS*, *CRYAA*, and *KCNE1* when mapping to the masked GRCh38.

We worked with the Genome Reference Consortium to use a new masking file that changes the sequence in the falsely duplicated regions of chromosome 21 on GRCh38 to N’s. Masking in this way maintains the same coordinates but dramatically improves variant calling in the genes. Previous work demonstrated that variant calls could be recovered even from short reads by masking extra copies of highly homologous “camouflaged” gene sequences, though this approach does not determine in which gene copy the variants occurred^8,28^. In our case, we are masking extra gene copies that are incorrect in the reference, enabling unambiguous variant calling in the correct genes. We show that masking the false duplications substantially improves recall and precision of variant calls in these genes for Illumina, PacBio HiFi, and ONT mapping-based methods, increasing sensitivity of Illumina-bwamem-GATK from 8% to 100% (**Figure 4**), without increasing errors in other regions (**Supplementary Figure 8**).

Our new benchmark also identified some falsely duplicated genes in GRCh37, specifically the medically relevant genes *MRC1* and *CNR2*. Both short and long reads map correctly to *MRC1* in GRCh38, but many reads incorrectly align to a false extra copy of the gene in GRCh37. Similarly, *CNR2* is annotated on GRCh37 to include a large region downstream that has an erroneous extra unplaced contig on chromosome 1 (chr1_gl000191_random) that interferes with mapping to a 106 kb region that includes part of *CNR2* as well as other genes (*PNRC2* and *SRSF10*) not included in our medical gene list. Our benchmark correctly resolves all of these genes on both GRCh37 and GRCh38 because the assembled contigs align correctly for each haplotype.

The CMRG benchmark also identified false positives that were eliminated by adding hs37d5 decoy sequence to GRCh37, but also identified false negatives caused by the decoy. The hs37d5 decoy was created from assembled sequences not in the GRCh37 reference, and was used in Phase 2 of the 1000 Genomes Project to remove some false positives due to mismapped reads from these sequences^29^. In order to evaluate the impact of the decoy on variant call accuracy in our challenging medically relevant genes, we benchmarked HG002 Illumina-bwamem-GATK calls against the new benchmark with and without adding the hs37d5 decoy sequence to the GRCh37 reference. Using the decoy eliminated 1,272 false positive SNVs and INDELs in the medical gene benchmark, including 1,191 in *KMT2C*, 15 in *MUC5B*, and the remainder in clusters of false positives in LINEs, SINEs, and LTRs in other genes. However, using the decoy sequence also caused 78 SNV and INDEL false negatives, notably 52 in *CYP4F12* and 18 in *LMF1* due to falsely duplicating parts of these genes. Therefore, while the hs37d5 decoy improves overall performance of variant calling, it can cause some false negatives in important genes similar to the false duplications in the primary assemblies discussed above. A potential solution may be to mask the falsely duplicated portions of the hs37d5 decoy similar to the masking of false duplications in GRCh38.

### Benchmark reliably identifies variant calling errors in challenging genes

We evaluated the CMRG small variant benchmark by comparing 7 VCFs from short- and long-read technologies and a variety of mapping and assembly-based variant calling methods. The goal of this curation process is to verify that the CMRG benchmark reliably identifies false positives and false negatives across sequencing technologies and variant calling methods. Manual curation of a random subset of 20 false positives, 20 false negatives, and 20 genotyping errors from each callset (split evenly between GRCh37 and GRCh38, and between SNVs and INDELs) demonstrated that most types of discrepancies were errors in each callset (**Supplementary Figure 9**). However, the majority of INDEL differences were identified as errors in the benchmark for two callsets, and curation identified 215 small regions with errors in the benchmark. These errors included missing haplotypes (particularly heterozygous INDELs in otherwise homozygous regions), as well as errors due to noise in the HiFi data in very long homopolymers, as detailed in **Supplementary Note 1**. We also excluded 33 errors found in manual curation of complex small variants in tandem repeats (**Supplementary Note 1**), such as *MUC5B* in **Supplementary Figure 10**. To more completely exclude these errors in the CMRG benchmark, we also curated all of the false positives, false negatives, and genotyping errors that were in at least half of the callsets on GRCh37 or GRCh38. We found that 44/50 and 59/63 of the errors identified by the evaluation on GRCh37 and GRCh38, respectively, were excluded by curation of the common false positives and false negatives. After excluding these errors, v1.00 accurately identifies errors for both SNVs and INDELs. We’ve included our full curation results in **Supplementary File 5**, which gives coordinates of common errors on both GRCh37 and GRCh38. This table can be used as a resource for investigating false positives and false negatives identified in a user’s query callset, since we provide notes about the evidence for the benchmark at each common false positive or false negative site.

We evaluated the CMRG SV benchmark by comparing four short and long read-based callsets, finding the benchmark reliably identified false positives and false negatives across all 4 callsets. Upon manual curation only two sites were identified as problematic due to different representations that current benchmarking tools could not reconcile. We also found the benchmarking statistics were sensitive to benchmarking tool parameters, particularly for duplications (**Supplementary Note 1**). We also compared Bionano optical mapping-based SV calls to the 50 benchmark SVs >=500 in size. Because many of these SVs were near the limit of detection of optical mapping, we curated these calls, and all were supported by the Bionano data.

From the manual curation of common false positives, false negatives, and genotyping errors, we also identified some categories of variants where the benchmark correctly identified errors in the majority of callsets: (1) clusters of false negatives and genotyping errors in the genes that are falsely duplicated in GRCh37 (*MRC1* and part of *CNR2*) and GRCh38 (*CBS*, *CRYAA*, *KCNE1*, and *H19*); (2) clusters of false positives and genotyping errors due to mismapped reads in the parts of *KMT2C* that are duplicated in HG002 relative to GRCh37 and GRCh38, which are responsible for 277 of the 386 false positives in the HiFi-DeepVariant callset (**Supplementary Figure 11**). We also determined that the benchmark correctly identified false negatives across technologies, but particularly short read-based methods, in segmental duplications like *SMN1* and *NCF1*, and in gene conversion-like events in *RHCE*, *SIGLEC16*, and *GTF2IRD2*. In addition to previously developed stratifications for difficult regions, we developed new stratifications for falsely duplicated genes, genes with large duplications, and complex variants in tandem repeats (**Supplementary Note 2**).

We further confirmed 225 of 226 variants in 10 genes in segmental duplications that were covered confidently by an orthogonal long range PCR and Sanger sequencing method (**Supplementary Table 1 and Supplementary File 6**). 127 other variants we attempted to confirm did not have coverage or had noisy sequencing, and only one variant (a homozygous SNV at GRCh38 chr16:2113578 in *PKD1*) was contradicted by long range PCR but was clearly supported by Illumina, 10x Genomics, PacBio HiFi, and ONT (**Supplementary Figure 12).**

To demonstrate how the CMRG benchmark can identify new types of errors relative to v4.2.1, we benchmarked a stringently-filtered Illumina-bwamem-GATK callset vs. both the v4.2.1 benchmark and the new medically relevant gene benchmark. **Figure 5** shows that the fraction not assessed decreases and the false negative rate increases substantially overall, but particularly for difficult variants. For SNVs, these difficult variants fall primarily in segmental duplications and low mappability regions, while for INDELs the CMRG also identifies additional false negatives in other regions excluded from the “Not in All Difficult” stratification, such as tandem repeats and homopolymers.

**Figure 5:**
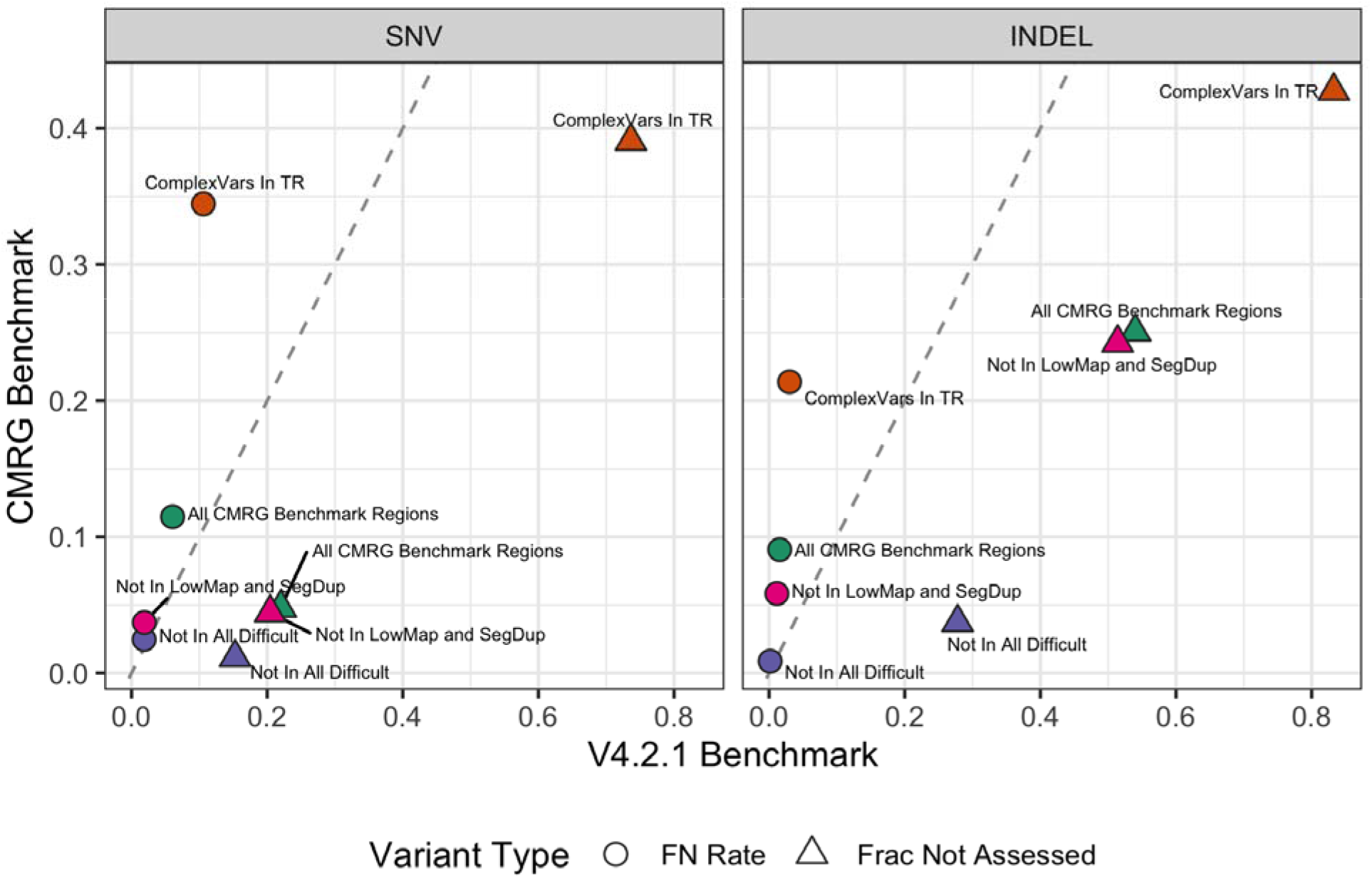
The new CMRG small variant benchmark includes more challenging variants and identifies more false negatives in a standard short-read callset (Illumina-bwamem-GATK) than the previous v4.2.1 benchmark in these challenging genes. While the false negative rate (circles) is similar in easier regions (purple “Not In All Difficult” points), the false negative rate is much higher overall (green “All CMRG Benchmark Regions” points). The fraction of variants excluded from the benchmark regions (triangles) is much higher for the v4.2.1 benchmark in all stratifications. This information is also presented in “summary stats NYGC” in **Supplementary File 4.**

### Remaining challenges across medically relevant genes

While the CMRG benchmark covers many new, challenging genes, 122 autosomal genes covered <90% by v4.2.1 are still excluded from the CMRG benchmark (110 on GRCh37 and 100 on GRCh38) for multiple reasons detailed in Supplementary File 7: When progressively categorizing excluded genes on GRCh38, (1) 20 genes were affected by gaps in the reference, (2) 38 genes had evidence of duplications in HG002 relative to GRCh38, (3) 6 genes were resolved, but excluded due to being in the MHC^14^, (4) 3 genes were resolved on GRCh38 but not GRCh37, as we required genes to be resolved on both references, (5) 19 were >90% included by the dip.bed but had multiple contigs or a break in the assembly-assembly alignment, (6) 7 had a large deletion of part or all of the gene on one haplotype, (7) 4 had breaks or false duplications in the hifiasm assembly, (8) 2 were in the structurally variable immunoglobulin locus, and (9) one (*TNNT3*) had a structural error in GRCh38 described in ^27^.

As examples, *LPA* and *CR1* were not included in the benchmark due to very large insertions and deletions, respectively, that cause a break in contig alignments, though the hifiasm assembly resolved both haplotypes (**Supplementary Figure 13 and 14)**. *LPA* contains multiple tandemly duplicated copies of the same region (i.e., kringle IV repeats with a unit length of ~5550 bp) that are associated with cardiovascular disease risk and is thus important to resolve correctly^30^. The HG002 hifiasm assembly resolved the entire *LPA* region, and the 44.1 kb and 99.9 kb expansions of the kringle IV repeats for the maternal and paternal haplotypes, respectively, were consistent with the insertions predicted by an independent trio-phased Bionano optical mapping assembly (45.0 kb and 101.2 kb). This complex, large expansion of the kringle IV repeats can be represented in many different ways in a VCF with different levels of precision (e.g., as a large insertion, a tandem duplication, or a CNV, and the copies may differ or include small variants). Existing benchmarking tools cannot compare these different representations robustly, partly limited by the VCF format^31^. To benchmark assemblies of this gene in HG002, the sequences could be compared directly to the hifiasm contigs, which we have annotated for *LPA* and other genes using LiftOff^32^. *CR1*, a gene implicated in Alzheimer’s disease^8^, is similarly resolved by hifiasm, containing a 18.5 kb homozygous deletion consistent with Bionano, but it causes a break in the dipcall/minimap2 alignment (**Supplementary Figure 14)**.

Other genes are excluded from the benchmark because they have extra copies in HG002 but not in GRCh38. For example, KCNJ18 is excluded because GRCh37 and GRCh38 are missing a copy of this gene (KCNJ17), so extra contigs from KCNJ17 align to KCNJ18.^27^ Also, genes in the KIR region are highly variable and CNVs are observed frequently in the population, with 35 alternate loci and 15 novel patches in GRCh38.p13. Hifiasm resolves the paternal allele in a single contig, but the maternal allele is split into 3 contigs in the KIR region, including a tandem duplication of the gene *KIR2DL1* (**Supplementary Figure 15**). There is no standard way to represent or benchmark small variants within duplicated regions, so we excluded *KCNJ18*, *KIR2DL1,* and other duplicated genes like *PRSS1* and *DUX4* from our benchmarks (**Supplementary Figures 16 and 17**). More information about these complex genes is in **Supplementary Note 3**.

## Discussion

In this work, we provide highly curated benchmarks for both phased small and structural variants covering 273 medically relevant and challenging genes. Parts or all of these genes are often excluded from standard targeted sequencing and whole-genome sequence (WGS) analysis as they include complex repetitive elements or are highly polymorphic. Still, the impact of these genes are well documented across multiple diseases and multiple studies. Our newly developed benchmark will help enable development and optimization of new methods to include them in genomic analysis. Thus, it paves the way to obtain comprehensive insights in these highly important regions of the human genome to further expand medical diagnoses and potentially improve understanding of the heritability for multiple diseases^33^. We give specific examples of challenges with calling variants in these genes, including mapping challenges for different technologies and identifying genes for which GRCh37 or GRCh38 is a better reference. This new benchmark was designed to be complementary to previous mapping-based benchmarks. Some difficult genes like *PMS2* are resolved well by v4.2.1 in HG002 and not by the assembly. Some difficult genes like the HLA family^14^ or *GBA1*/*GBA2* are resolved well by the assembly, but are not included in the new benchmark because they were well-resolved previously.

Still, a few challenging regions remain excluded from our benchmark or are not resolvable despite the availability of highly accurate long read data. Some genes include variable long tandem repeats (e.g. *LPA* and *CR1*), which are resolved in our assembly, but the large >20 kb changes in length of the alleles are currently too complex for standard benchmarking methodologies. This clearly shows the need for more advanced methods, potentially graph representations of haplotypes or alleles. In addition, a few genes (e.g., *SMN2*) escaped a comprehensive and accurate assessment even with current long read-based assembly methods, highlighting the need for further development of sequencing and bioinformatics methods. Furthermore, our extensive curation of the benchmark helped identify limitations of the current whole genome diploid assembly methods, paving the way for future whole genome assembly-based benchmarks: (1) The assembly often misses one allele for heterozygous INDELs in highly homozygous regions. (2) Some consensus errors exist, causing errors in a single read to be called as variants. (3) If both haplotypes of the assembly do not completely traverse segmental duplications, the assembly is less reliable (e.g., *SMN2* in HG002) though it sometimes can still be correct (e.g., *SMN1* in HG002). Some genes also may be resolvable in HG002 but not in other genomes or vice versa, due to structural or copy number variability in the population, so benchmarks for additional samples will be needed.

By basing this benchmark on a whole genome diploid assembly, we are able to identify biases in mapping-based methods due to errors in the GRCh37 and GRCh38 references. While previous studies concluded that variant calling performance is generally better on GRCh38^34,35^, our benchmark demonstrates that variant calls in some genes are less accurate on GRCh38 than GRCh37. However, we demonstrate that masking false duplications on GRCh38 greatly improves performance in these genes. Interestingly, another group recently independently identified the importance of masking the extra copy of one gene (*U2AF1/U2AF1L5*) for cancer research^36^. Our results identify that false duplications cause many of the discrepancies found recently between exome variant calls on GRCh37 and GRCh38,^37^ and highlight the importance of our proposed masked GRCh38 genome. Previous benchmarks excluded regions that were problematic for each reference. By aligning the HG002 assembly to each reference, we produced similar benchmarks for both versions of the reference, so that users can better understand strengths and weaknesses of each reference, and test modifications to the reference such as the hs37d5 decoy for GRCh37 or the masked GRCh38 we propose here. During this process, we also identified and resolved variant calling errors due to several false duplications in these medically important genes in GRCh38 on chromosome 21. Overall, 11 genes are impacted by these false duplications, including 3 medically relevant genes from our list: *CBS*, *KCNE1* and *CRYAA*. As a solution to this we provide a new GRCh38 reference that masks the erroneous copy of the duplicated genes. We use our benchmark to show this reference dramatically improves read mapping and variant calling in these genes across almost all sequencing technologies. These false duplications exist only in GRCh38 and not in other human reference genome versions or in the broader population. A new telomere-to-telomere reference genome eliminates these false duplications and fixes collapsed duplications that prevented us from creating a benchmark for medically relevant genes like *KCNJ18* and *MAP2K3*, and a similar CMRG benchmark for HG002 is now available on the new reference^27^. Future work will include using phased, diploid assemblies to form benchmarks for more genic and non-genic regions of the genome, eventually using genomes that are assembled telomere-to-telomere.

This benchmark is additionally unique as it combines the small and structural variant calls that have been so often held separate, despite the arbitrary 50 bp size cutoff between them^5^. The phased assembly contigs used in this work^2^ provide a natural way to combine SV and small variants. The combination of both allele types is an important step towards a more comprehensive benchmark that takes into account the relationship between nearby small and structural variants. Thus, this benchmark indeed represents a comprehensive snap shot of HG002 as a representative for these 273 medically relevant regions. Furthermore, this promotes the trend of SVs and phasing of complex variants becoming more widely used for medical research and clinical sequencing^5,38^. While the polymorphisms that we identified here are likely benign, it is clear that this benchmark will improve methods to assess these regions and maybe more importantly enable geneticists to assess the reliability of their methods or pipelines at hand.

Our new approach to form benchmarks from a phased whole genome assembly is a prototype for future comprehensive benchmarks covering the whole genome. Overall, this benchmark enables a more comprehensive assessment of sequencing strategies, analytical methodologies and other developments for challenging genomic variants and regions important to medical research, paving the way for improved clinical diagnoses.

## Methods

### Sample availability

For the 10× Genomics and Oxford Nanopore sequencing and Bionano mapping, the GM24385 (RRID:CVCL_1C78) cell line was obtained from the Coriell Institute for Medical Research National Institute for General Medical Sciences cell line repository. For the Illumina and Pacific Biosciences sequencing, NIST RM 8391 DNA was used, which was prepared from a large batch of GM24385 to control for differences arising during cell growth. For binning reads into paternal and maternal haplotypes, Illumina sequencing of DNA from NIST RM 8392 (HG002-HG004) was used. DNA was extracted from cell lines publicly available as GM24149 (RRID:CVCL_1C54) and GM24143 (RRID:CVCL_1C48) at the Coriell Institute for Medical Research National Institute for General Medical Sciences cell line repository.

### Medical Genes

We used genes from a variety of databases and sources in order to compile a list of medically-relevant genes. The largest set of genes we use is from Mandelker *et al*. Supplementary Table 13 which was a capture of the OMIM, HGMD, and ClinVar databases gathered around 2012. Further, we used the COSMIC cancer gene census which is a list of 723 genes. **Supplementary File 1** also contains additional details about the higher priority list of 942 genes in the union of ClinGen genes with “definitive”, “strong” or “moderate” evidence (719 genes), NCCN/ESMO (hereditary cancer syndromes) (49 genes), ACMG SF 2.0 (commonly referred to as the ACMG59 - for which reporting of secondary or incidental findings are recommended) (59 genes), CPIC pharmacogenetics genes (127 genes), and the Counsyl expanded carrier screening list (235 genes), which includes recommended reproductive medicine genes as a small subset.

### Medical Gene Coordinate Discovery

We used coordinates from ENSEMBL (https://uswest.ensembl.org/index.html) then downloaded “chromosome”, “start”, “end”, “gene_name”, and “stable_ID” using bioMart for GRCh38 and GRCh37. We looked up the collection of medical genes and found the coordinates for each in GRCh38 and GRCh37, available under https://github.com/usnistgov/cmrg-benchmarkset-manuscript/tree/master/data/gene_coords/unsorted.

### Calculating Overlap with GIAB HG002 v4.2.1 small variant benchmark

We used bedtools^39^ intersect with the ENSEMBL coordinates for each gene and the v4.2.1 small variant benchmark regions BED. We calculated the number of bases in the intersection and compared that to the total number of bases in each gene. We chose 90% as the threshold for the purpose of keeping manual curation tractable over the set of the genes.

### Diploid Assembly using PacBio HiFi reads with hifiasm using trio-binning

We use the diploid assembly produced by hifiasm v0.11 using 34x coverage (two 15 kb and two 20 kb libraries) by PacBio HiFi Sequel II System with Chemistry 2.0 reads (https://github.com/human-pangenomics/HG002_Data_Freeze_v1.0#hg002-data-freeze-v10-recommended-downsampled-data-mix) using kmer information from parental Illumina short reads (30x 2×150bp reads at https://s3-us-west-2.amazonaws.com/human-pangenomics/index.html?prefix=NHGRI_UCSC_panel/HG002/hpp_HG002_NA24385_son_v1/parents/ILMN/downsampled/), described recently^2^.

### Calling variants relative to GRCh37 and GRCh38 using dipcall

We aligned the diploid assembly of HG002 to GRCh37 and GRCh38 using minimap2^40^ through dipcall (https://github.com/lh3/dipcall) as is done in the NIST assembly benchmarking pipeline (https://github.com/usnistgov/giab-asm-benchmarking). Dipcall generates variant calls using any non-reference support in regions that are greater than or equal to 50 kb with contigs having mapping quality greater than or equal to 5. Dipcall also produces a BED which denotes confident regions that are covered by an alignment greater than or equal to 50 kb with contigs having mapping quality greater than or equal to 5 and with no other greater than 10 kb alignments.

### Benchmark Development

We selected genes that had continuous haplotype coverage of the gene body including the 20 kb on each side to account for robust alignments. In addition, each haplotype had to fully cover any segmental duplications in close proximity or overlapping the extended gene regions. This also included complex SVs inside of the segmental duplications to be able to robustly identify SNVs and SVs subsequently. We considered a gene to be fully resolved by the diploid assembly if the dip.bed covered the gene along with 20 kb of flanking sequence to consider the PacBio HiFi read length as well as any overlapping segmental duplications. We chose these criteria to ensure that genes were resolved in regions with high-quality assembly.

We then performed manual curation of the resolved genes and flanking sequence to understand overall characteristics of the new candidate benchmark. We began initial evaluation against mapping-based callsets to understand the performance of the benchmark in these genes. We identified that perfect homopolymers > 20 bp and imperfect homopolymers > 20 bp accounted for a majority of false negatives and false positives for both SNPs and INDELs. Imperfect homopolymers are defined as stretches of one base that are interrupted by one different base in one or more locations, and each of the stretches of exact homopolymer bases have to be at least 4 bp (e.g., AAAAGAAAAAGAAAATAAAA). Manual curation of a random subset of these sites showed that in most instances it was unclear whether the mapping-based callset or the assembly-based benchmark was correct. Bed files for these homopolymers are available under https://ftp-trace.ncbi.nlm.nih.gov/ReferenceSamples/giab/release/genome-stratifications/.

We exclude the following regions from the v0.02.03 small variant and v0.01 structural variant benchmark regions (the benchmark versions used in the evaluation): 1) one region identified manually as an erroneous insertion resulting from an issue with the method hifiasm v0.11 used to generate the consensus sequence, 2) genes in the MHC, since these were previously resolved by diploid assembly in the v4.2.1 benchmark^14^, and 3) regions around variants identified as errors or unclear upon manual curation, as described below. For the small variant benchmark, we additionally exclude: 1) structural variants at least 50 bp in size and overlapping tandem repeats because these cannot be compared robustly with small variant comparison tools, and 2) perfect and imperfect homopolymers > 20 bp + 5 bp on each side. For the structural variant benchmark, we additionally exclude 1) tandem repeats that contain more than one variant at least 10 bp in size because these complex variants can cause inaccurate comparisons with current benchmarking tools, and 2) INDELs 35 bp to 49 bp in size.

### Benchmark Evaluation

We used hap.py^41^ with vcfeval to compare VCFs from a variety of sequencing technologies and variant calling methods to the GRCh37 and GRCh38 difficult medical gene small variant benchmark, with GIAB/GA4GH stratifications under https://ftp-trace.ncbi.nlm.nih.gov/ReferenceSamples/giab/release/genome-stratifications/. We randomly selected 60 total sites for curation, with 30 selected from GRCh37 and 30 from GRCh38. 5 SNVs and 5 INDELs were selected from each of these 3 categories: (1) False Positives – variant in comparison VCF but not in Benchmark, (2) False Negative – variant not in comparison VCF but in Benchmark, and (3) Genotype errors - variant appears as both a False Positive and a False Negative using hap.py with vcfeval. This curation process will also help us to make further refinements, if needed, to the GIAB benchmark. For the small variant benchmark evaluation, we used 7 VCFs^12^ from short- and long-read technologies and a variety of mapping and assembly-based variant calling methods: 1) Illumina-DRAGEN, 2) Illumina-NovaSeq-GATK4^42^, 3) Illumina-xAtlas^43^, 4) PacBio HiFi-GATK4, 5) an assembly based on ONT reads called with dipcall, 6) union of three callsets: Illumina called with modified GATK, PacBio HiFi called with Longshot^44^ v0.4.1, and ONT called with PEPPER-DeepVariant^45^, and 7) Illumina + PacBio + ONT combined called with NeuSomatic^46^. We exclude errors identified upon curation, as described in Supplementary Note 1. In Supplementary Note 2, we also include performance metrics for the 53x 15 kb + 20 kb HiFi-DeepVariant v0.9 callset under https://ftp-trace.ncbi.nlm.nih.gov/ReferenceSamples/giab/data/AshkenazimTrio/analysis/PacBio_CCS_15kb_20kb_chemistry2_10312019/

### Variant Callsets Used for Evaluation

#### NCBI de novo assembly

The de novo assembly of HG002 was initially generated using NextDenovo with ONT Promethion data (ftp://ftp-trace.ncbi.nlm.nih.gov/giab/ftp/data/AshkenazimTrio/HG002_NA24385_son/UCSC_Ultralong_OxfordNanopore_Promethion/), then polished with PacBio 15/20 kb CCS reads (ftp://ftp-trace.ncbi.nlm.nih.gov/giab/ftp/data/AshkenazimTrio/HG002_NA24385_son/PacBio_CCS_15kb_20kb_chemistry2/reads/), followed by scaffolding with HiC data (https://github.com/human-pangenomics/HG002_Data_Freeze_v1.0). The scaffolded assembly was further polished with Illumina short reads (https://github.com/human-pangenomics/HG002_Data_Freeze_v1.0) twice using pilon^47^, and then phased with Whatshap^48^. Finally, two vcf files (HG002_grch37_dipcall.vcf.gz and HG002_grch38_dipcall.vcf.gz) were generated based on phased HG002 genome using dipcall with GRCh37 and GRCh38 reference genomes, respectively (ftp://ftp-trace.ncbi.nlm.nih.gov/giab/ftp/data/AshkenazimTrio/analysis/NCBI_variant_callsets_for_medicalgenes_evaluation_10282020/).

#### Illumina-DRAGEN

HG002 DNA was prepared using Illumina DNA PCR-Free Library Prep. The library was sequenced on NovaSeq 6000 with 151 bp paired-end reads. Illumina DRAGEN 3.6.3 was used to align sequencing reads and call variants. SNP and INDEL were filtered using the following hard filters:

DRAGENHardSNP:snp: MQ < 30.0 || MQRankSum < −12.5 || ReadPosRankSum < - 8.0;DRAGENHardINDEL:indel: ReadPosRankSum < −20.0

#### Illumina + PacBio + ONT combined called with NeuSomatic

The predictions are based on the adaptation of the deep learning based framework in NeuSomatic for germline variant calling. We used the network model trained for NeuSomatic’s submission for the PrecisionFDA truth challenge v2^12^. The model is trained on HG002 using GIAB benchmark set V4.2. For this callset separate input channels were used for PacBio, Illumina, and ONT reads.

#### DNAnexus - Union of short read callsets from 4 callers

We downloaded HG002 WGS FASTQ reads from NIST’s FTP ([ftp://ftp-trace.ncbi.nlm.nih.gov/giab/ftp/data/AshkenazimTrio/HG002_NA24385_son/NIST_Illumina_2x250bps/reads/](ftp://ftp-trace.ncbi.nlm.nih.gov/giab/ftp/data/AshkenazimTrio/HG002_NA24385_son/NIST_Illumina_2x250bps/reads/))^49^, followed by downsampling the reads to 35X coverage (47.52%) from originally 73.65X coverage using ‘seqtk’.

We called variants against both GRCh38 (GRCh38 primary contigs + decoy contigs, but no ALT contigs nor HLA genes) and hs37d5 builds. We run four different germline variant callers (see below) with their suggested default parameters, collected the union of all variants using a customized script where we recorded which caller(s) called the variant and their filter statuses in INFO and FILTER fields, and generated the union VCF file for HG002. The customized script excludes variants (with the same CHROM, POS, REF, ALT) that have conflicting genotype (GT) reported by different callers, and only keeps the variants that are reported as exactly the same genotype when more than one caller is calling it. The variant calling pipelines used were: (1) BWA-MEM+GATK4 ([BWA-MEM^50^ version 0.7.17-r1188] (https://github.com/lh3/bwa) + [GATK version gatk-4.1.4.1] (https://gatk.broadinstitute.org/hc/en-us)); (2) Parabricks_DeepVariant ([Parabricks Pipelines DeepVariant v3.0.0_2](https://developer.nvidia.com/clara-parabricks)); Sentieon_DNAscope | [Sentieon (DNAscope) version sentieon_release_201911] (https://www.sentieon.com/products/#dnaseq)); (4) BWA-MEM+Strelka2 ([BWA-MEM version 0.7.17-r1188](https://github.com/lh3/bwa) + [Strelka2 version 2.9.10](https://github.com/Illumina/strelka).

#### Illumina Novaseq 2×250bp data

The sample HG002 was sequenced on an Illumina Novaseq 6000 instrument with 2×250bp paired end reads at the New York Genome Center. The libraries were prepped using TruSeq DNA PCR-free library preparation kit. The raw reads were aligned to both GRCh37 and GRCh38 human reference. Alignment to GRCh38 reference, marking duplicates and base quality recalibration was performed as outlined in the Centers for Common Disease Genomics (CCDG) functional equivalence paper^51^. Alignment to GRCh37 was performed using BWA-MEM ^50^(ver. 0.7.8) and marking duplicates using Picard (ver. 1.83) and local INDEL realignment and base quality recalibration using GATK^52^ (ver. 3.4-0). Variant calling was performed using GATK (ver. 3.5) adhering to the best practices recommendations from the GATK team. Variant calling constituted generating gVCF using HaplotypeCaller, genotyping using the GenotypeGVCFs subcommand and variant filtering performed using VariantRecalibrator and ApplyRecalibration steps. A tranche cutoff of 99.8 was applied to SNP calls and 99.0 to INDELs to determine PASS variants. The raw reads are available for download at SRA at https://www.ncbi.nlm.nih.gov/sra/SRX7925517

#### Small variants from Illumina + PacBio + ONT

For this submission we combined three sequencing technologies data to obtain a more sensitive VCF file.

We used our in-house variant calling pipeline for the Illumina dataset. In short, BWA-MEM v0.7.15-r1140 was used to align reads to GRCh37 or GRCh38 reference genome and BAM files were processed with SAMtools^53^ v1.3 and Picard v2.10.10. SNVs and INDELs were identified with the HaplotypeCaller following the Best Practices workflow recommendations for germline variant calling in GATK v3.8. ^50^

For both PacBio and ONT datasets, we run another pipeline using NanoPlot v1.27.0 for quality control, Filtlong v0.2.0 for filtering reads, and minimap2 v2.17-r941 for alignment. Longshot v0.4.1 was used for variant calling for PacBio data and PEPPER-DeepVariant for ONT data.

On the variant callsets, we filtered out variants applying the following criteria: FILTER=PASS, QD (*Quality by Depth*)≥2.0 and MQ (*Mapping Quality*)≥50 for Illumina data; and FILTER=PASS and QUAL (*Quality*)≥150 for PacBio data. No filters were applied to ONT calls.

Finally, we created a consensus VCF file by merging the single VCF files obtained by each of these three pipelines using the GATK CombineVariants tool.

#### *Structural variants from* Illumina (intersection callsets from 5 callers)

We called SVs on short-read Illumina data using 5 different SV callers: DELLY^54^ v0.8.5, GRIDSS^55^ v2.9.4, LUMPY^56^ v0.3.1, Manta^57^ v1.6.0, and Wham^58^ v1.7.0. The HG002 BAM file aligned to GRCh37 or GRCh38 reference genome by BWA-MEM v0.7.15-r1140, with duplicates marked using Picard v2.10.10, and base quality score recalibrated by GATK v3.8, was used to feed these SV callers, which were executed with recommended default parameters. LUMPY and Wham SV calls were genotyped using SVTyper v0.7.1. GRIDSS SV types were assigned with the *simple-event-annotation* R script, included in the GRIDSS package.

Resulting SV callsets were filtered based on author’s recommendations for each caller: Manta (FILTER=PASS, INFO/PRECISE, FORMAT/PR ≥10), LUMPY (INFO/PRECISE, remove genotypes 0/0, QUAL ≥100, FORMAT/AO ≥7), DELLY (FILTER=PASS, INFO/PRECISE), GRIDSS (FILTER=PASS, INFO/PRECISE, QUAL ≥1,000, INFO/SVLEN <1 kb, remove DAC Encode regions), and Wham (INFO/SVLEN <2kb, INFO/A >5, remove genotypes 0/0, INFO/CW[bnd] >0.2).

The resulting VCF files from each caller were merged to create the intersection of variants using SURVIVOR^59^ v1.0.7, containing variants >50 bp in size, with 1,000 bp as the distance parameter and without requiring any type specificity (all variant types are merged). In the intersection set, we retained calls supported by two or more callers.

#### Structural variants from ONT (merge callsets from 2 callers)

For these submissions, we built a custom pipeline to process the ONT HG002 dataset using NanoPlot^60^ v1.27.0 for quality control, Filtlong v0.2.0 for filtering reads, and minimap2^40,60^ v2.17-r941 for alignment to the GRCh37 or GRCh38 reference genome.

The structural variants were called on the resulting BAM file using cuteSV v1.0.8 and Sniffles^61^ v1.0.12. The resulting VCF files were filtered out based on the default values suggested by each tool authors: cuteSV ^62^ (minimum read support of 10 reads, INFO/RE ≥10; this caller intrinsically filters by FILTER=PASS and INFO/PRECISE) and Sniffles (FILTER=PASS, INFO/PRECISE and minimum read support of 10 reads).

Finally, we created the VCF files by merging the single filtered VCF files using SURVIVOR ^59^ v1.0.7, containing variants >50 bp in size, with 1,000 bp as the distance parameter and without requiring any type specificity (all variant types are merged).

### Remapping Variants Between GRCh38 and GRCh37

To remap curated variant locations between GRCh38 and GRCh37, we used the NCBI Remap tool. For variants that remapped in the first pass, we used the first pass location. For variants that did not remap in the first pass, all remapped in the second pass, and we used the second pass location.

### Masking false duplications on chromosome 21 of GRCh38

We worked with the Genome Reference Consortium (GRC) to develop a list of regions in GRCh38 that could be masked without changing coordinates or harming variant calling, because they were erroneously duplicated sequences or contaminations. The BED file with these regions at https://ftp.ncbi.nlm.nih.gov/genomes/all/GCA/000/001/405/GCA_000001405.15_GRCh38/seqs_for_alignment_pipelines.ucsc_ids/GCA_000001405.15_GRCh38_GRC_exclusions.bed. To create the masked reference, we started with the GRCh38 reference with no ALT loci nor decoy from ftp://ftp.ncbi.nlm.nih.gov/genomes/all/GCA/000/001/405/GCA_000001405.15_GRCh38/seqs_for_alignment_pipelines.ucsc_ids/GCA_000001405.15_GRCh38_no_alt_analysis_set.fna.gz.

To generate the masked GRCh38 (i.e., replacing the duplicated and contaminated reference sequence with N’s), we used the Bedtools tools (https://github.com/arq5x/bedtools2) command: maskFastaFromBed -fi GCA_000001405.15_GRCh38_no_alt_analysis_set.fasta -bed GCA_000001405.15_GRCh38_GRC_exclusions.bed -fo GCA_000001405.15_GRCh38_no_alt_analysis_set_maskedGRC_exclusions.fasta

To generate the v2 masked GRCh38, we ran the Bedtools tools (https://github.com/arq5x/bedtools2) command:

‘‘‘

maskFastaFromBed -fi GCA_000001405.15_GRCh38_no_alt_analysis_set.fasta -bed GCA_000001405.15_GRCh38_GRC_exclusions_T2Tv2.bed -fo GCA_000001405.15_GRCh38_no_alt_analysis_set_maskedGRC_exclusions.fasta

This uses the a bed file GCA_000001405.15_GRCh38_GRC_exclusionsv2.bed generated by the Telomere to Telomere Consortium Variants team to mask false duplications located under https://ftp-trace.ncbi.nlm.nih.gov/ReferenceSamples/giab/release/references, which also contains the new masked references and other references used in this work.

### Evaluation of GRCh38 masked genome improvement

For short reads, a common whole genome resequencing analysis pipeline was used to produce variant call files for the HG002 sample in VCF and gVCF formats. The applications and parameters used in the analysis pipeline were derived from best practices for Illumina short-read WGS resequencing analysis developed for the Centers for Common Disease Genomics (CCDG) project^51^. The analysis pipeline consists of the following high-level steps:

1. Sequence alignment to reference genome using BWA-MEM
2. Duplicate read marking using Picard Tools MarkDuplicates
3. Base quality score recalibration using GATK BaseRecalibrator
4. Variant calling using GATK HaplotypeCaller

This analysis pipeline was run twice on a set of paired-end 35x HG002 FASTQs as input, with the pipeline runs differing only by reference genome used during the alignment step. The first run used a version of the GRCh38 reference genome prepared without decoy or alternate haplotype contigs. The second run used a version of the GRCh38 reference genome identical to that used in the first run, except that five regions in chr21 and the entire contig chrUn_KI270752v1 were masked with N’s, as described above.

The commands executed by the analysis pipeline runs are in Supplementary File 8 and Supplementary File 9, which correspond to the runs using the unmasked and masked GRCh38 references genomes, respectively.

The following versions were used for applications and resources used in the analysis pipeline: BWA v0.7.15, GATK v3.6, Java v1.8.0_74 (OpenJDK), Picard Tools v2.6.0, Sambamba^63^ v0.6.7, Samblaster^64^ v0.1.24, Samtools v1.9, dbSNP Build 138 on GRCh38, and Known INDELs from Mills and 1000 Genomes Project on GRCh38.

For PacBio long reads, we used a 35x 15 kb + 20 kb HiFi dataset from the precisionFDA Truth Challenge V2,^12^ aligned to the standard and masked GRCh38 reference with pbmm, and called variants with DeepVariant v1.0.^65^ **For PacBio lHiFi long readsMasked GRCh38 region**

### HG002 Diploid Annotation

Liftoff^32^ v1.4.0 was used with default parameters to lift over Ensembl v100 annotations from GRCh38 onto each haplotype assembly separately. The resulting gff files are available under https://ftp-trace.ncbi.nlm.nih.gov/ReferenceSamples/giab/release/AshkenazimTrio/HG002_NA24385_son/CMRG_v1.00/hifiasm-assembly/

## Code availability

Scripts used to develop the CMRG benchmark and generate figures and tables for the manuscript are being made available at https://github.com/usnistgov/cmrg-benchmarkset-manuscript and archived at https://doi.org/10.18434/mds2-2475. The previously developed assembly, which was used as the basis of this benchmark, was from hifiasm v0.11. A variety of open source software was used for variant calling for the evaluations of the benchmark: NextDenovo, DRAGEN 3.6.3, NeuSomatic’s submission for the PrecisionFDA truth challenge v2^12^, [BWA-MEM^50^ version 0.7.17-r1188] (https://github.com/lh3/bwa), [GATK version gatk-4.1.4.1] (https://gatk.broadinstitute.org/hc/en-us)); Parabricks_DeepVariant ([Parabricks Pipelines DeepVariant v3.0.0_2](https://developer.nvidia.com/clara-parabricks)); Sentieon (DNAscope) version sentieon_release_201911 (https://www.sentieon.com/products/#dnaseq)); BWA-MEM+Strelka2 ([BWA-MEM version 0.7.17-r1188](https://github.com/lh3/bwa) + [Strelka2 version 2.9.10](https://github.com/Illumina/strelka), BWA-MEM ^50^(ver. 0.7.8), Picard tools (https://broadinstitute.github.io/picard/) (ver. 1.83), GATK^52^ (ver. 3.4-0), GATK (ver. 3.5), BWA-MEM v0.7.15-r1140, SAMtools^53^ v1.3, Picard v2.10.10, GATK v3.8, DELLY^54^ v0.8.5, GRIDSS^55^ v2.9.4, LUMPY^56^ v0.3.1, Manta^57^ v1.6.0, and Wham^58^ v1.7.0, NanoPlot^60^ v1.27.0, Filtlong v0.2.0, minimap2^40,60^ v2.17-r941, cuteSV v1.0.8, Sniffles^61^ v1.0.12, SURVIVOR ^59^ v1.0.7, BWA v0.7.15, GATK v3.6, Java v1.8.0_74 (OpenJDK), Picard Tools v2.6.0, Sambamba^63^ v0.6.7, Samblaster^64^ v0.1.24, Samtools v1.9.

## Data availability

The v1.00 benchmark VCF and BED files, as well as Liftoff gene annotations, assembly-assembly alignments, and variant calls, are available at https://ftp-trace.ncbi.nlm.nih.gov/ReferenceSamples/giab/release/AshkenazimTrio/HG002_NA24385_son/CMRG_v1.00/ and archived at https://doi.org/10.18434/mds2-2475. This is released as a separate benchmark from v4.2.1 because it includes a small fraction of the genome, it has different characteristics from the mapping-based v4.2.1, and v4.2.1 only includes small variants. Using v4.2.1 and the CMRG benchmarks as two separate benchmarks enables users to obtain broader performance metrics for most of the genome and for a small set of particularly challenging genes, respectively. The masked GRCh38 reference, recently updated to version 2 with additional false duplications from the Telomere to Telomere Consortium, is under https://ftp-trace.ncbi.nlm.nih.gov/ReferenceSamples/giab/release/references. We recommend using v2.01 GA4GH/GIAB stratification bed files intended for use with hap.py when benchmarking, available under https://ftp-trace.ncbi.nlm.nih.gov/ReferenceSamples/giab/release/genome-stratifications/. These stratifications include bed files corresponding to false duplications and collapsed duplications in GRCh38. All data have no restrictions, as the HG002 sample has an open consent from the Personal Genome Project.

## Acknowledgments

We thank the Genome Reference Consortium for their curation efforts of GRCh37 and GRCh38 (https://www.genomereference.org), especially Valerie A Schneider and Paul A Kitts from NIH/NCBI for developing the falsely duplicated regions that should be masked in GRCh38. We thank Sierra Miller at NIST for helping make available benchmark sets and READMEs. Certain commercial equipment, instruments, or materials are identified to specify adequately experimental conditions or reported results. Such identification does not imply recommendation or endorsement by the National Institute of Standards and Technology, nor does it imply that the equipment, instruments, or materials identified are necessarily the best available for the purpose. CF was funded by Instituto de Salud Carlos III (PI20/00876) and Ministerio de Ciencia e Innovación (RTC-2017-6471-1; AEI/FEDER, UE), cofinanced by the European Regional Development Fund (ERDF) “A Way of Making Europe” from the European Union (EU), and Cabildo Insular de Tenerife (CGIEU0000219140). JMLS was funded by Consejería de Educación-Gobierno de Canarias and Cabildo Insular de Tenerife (BOC n.° 163, 24/08/2017). FJS and MM was supported by NIH (UM1 HG008898). Chunlin Xiao was supported by the Intramural Research Program of the National Library of Medicine, National Institutes of Health. KM was supported by NIH/NHGRI R01 1R01HG011274-01 and NIH/NHGRI U01 1U01HG010971. HL was supported by NIH (R01 HG010040 and U01 HG010961).

## References

1. Wenger, A. M. et al. Accurate circular consensus long-read sequencing improves variant detection and assembly of a human genome. Nat. Biotechnol. 37, 1155–1162 (2019).

2. Cheng, H., Concepcion, G. T., Feng, X., Zhang, H. & Li, H. Haplotype-resolved de novo assembly using phased assembly graphs with hifiasm. Nat. Methods 18, 170–175 (2021).

3. Nurk, S. et al. HiCanu: accurate assembly of segmental duplications, satellites, and allelic variants from high-fidelity long reads. Genome Res. 30, 1291–1305 (2020).

4. Shafin, K. et al. Nanopore sequencing and the Shasta toolkit enable efficient de novo assembly of eleven human genomes. Nat. Biotechnol. 38, 1044–1053 (2020).

5. Mahmoud, M. et al. Structural variant calling: the long and the short of it. Genome Biol. 20, 246 (2019).

6. De Coster, W., Weissensteiner, M. H. & Sedlazeck, F. J. Towards population-scale long-read sequencing. Nat. Rev. Genet. (2021) doi:10.1038/s41576-021-00367-3.

7. Mandelker, D. et al. Navigating highly homologous genes in a molecular diagnostic setting: a resource for clinical next-generation sequencing. Genet. Med. 18, 1282–1289 (2016).

8. Ebbert, M. T. W. et al. Systematic analysis of dark and camouflaged genes reveals disease-relevant genes hiding in plain sight. Genome Biol. 20, 1–23 (2019).

9. Lincoln, S. E. et al. One in seven pathogenic variants can be challenging to detect by NGS: An analysis of 450,000 patients with implications for clinical sensitivity and genetic test implementation. medRxiv 2020.07.22.20159434 (2020).

10. Zook, J. M. et al. An open resource for accurately benchmarking small variant and reference calls. Nat. Biotechnol. 37, 561–566 (2019).

11. Zook, J. M. et al. Author Correction: A robust benchmark for detection of germline large deletions and insertions. Nat. Biotechnol. 38, 1357 (2020).

12. Olson, N. D. et al. precisionFDA Truth Challenge V2: Calling variants from short- and long-reads in difficult-to-map regions. bioRxiv (2020) doi:10.1101/2020.11.13.380741.

13. Wagner, J. et al. Benchmarking challenging small variants with linked and long reads. bioRxiv (2020) doi:10.1101/2020.07.24.212712.

14. Chin, C.-S. et al. A diploid assembly-based benchmark for variants in the major histocompatibility complex. Nat. Commun. 11, 4794 (2020).

15. Goldfeder, R. L. et al. Medical implications of technical accuracy in genome sequencing. Genome Med. 8, 1–12 (2016).

16. Ball, M. P. et al. A public resource facilitating clinical use of genomes. Proc. Natl. Acad. Sci U. S. A. 109, 11920–11927 (2012).

17. Tate, J. G. et al. COSMIC: the Catalogue Of Somatic Mutations In Cancer. Nucleic Acids Res. 47, D941–D947 (2019).

18. Ross, M. G. et al. Characterizing and measuring bias in sequence data. Genome Biol. 14, R51 (2013).

19. Prior, T. W., Leach, M. E. & Finanger, E. Spinal Muscular Atrophy. in GeneReviews® [Internet] (University of Washington, Seattle, 2020).

20. Biros, I. & Forrest, S. Spinal muscular atrophy: untangling the knot? J. Med. Genet. 36, 1–8 (1999).

21. Leiding, J. W. & Holland, S. M. Chronic Granulomatous Disease. in GeneReviews® [Internet] (University of Washington, Seattle, 2016).

22. Innan, H. A two-locus gene conversion model with selection and its application to the human RHCE and RHD genes. Proc. Natl. Acad. Sci. U. S. A. 100, 8793–8798 (2003).

23. Hayakawa, T. et al. Coevolution of Siglec-11 and Siglec-16 via gene conversion in primates. BMC Evol. Biol. 17, (2017).

24. Garg, P. et al. Pervasive cis effects of variation in copy number of large tandem repeats on local DNA methylation and gene expression. Am. J. Hum. Genet. (2021) doi:10.1016/j.ajhg.2021.03.016.

25. Lennerz, J. K. et al. Addition of H19 ‘Loss of Methylation Testing’ for Beckwith-Wiedemann Syndrome (BWS) Increases the Diagnostic Yield. J. Mol. Diagn. 12, 576 (2010).

26. Nurk, S. et al. The complete sequence of a human genome. bioRxiv 2021.05.26.445798 (2021) doi:10.1101/2021.05.26.445798.

27. Aganezov, S. et al. A complete reference genome improves analysis of human genetic variation. bioRxiv 2021.07.12.452063 (2021) doi:10.1101/2021.07.12.452063.

28. Boisson, B. et al. Rescue of recurrent deep intronic mutation underlying cell type– dependent quantitative NEMO deficiency. Journal of Clinical Investigation vol. 129 583–597 (2018).

29. 1000 Genomes Project Consortium et al. A global reference for human genetic variation. Nature 526, 68–74 (2015).

30. Schmidt, K., Noureen, A., Kronenberg, F. & Utermann, G. Structure, function, and genetics of lipoprotein (a). Journal of Lipid Research vol. 57 1339–1359 (2016).

31. Li, H., Feng, X. & Chu, C. The design and construction of reference pangenome graphs with minigraph. Genome Biol. 21, 265 (2020).

32. Shumate, A. & Salzberg, S. L. Liftoff: accurate mapping of gene annotations. Bioinformatics (2020) doi:10.1093/bioinformatics/btaa1016.

33. Theunissen, F. et al. Structural Variants May Be a Source of Missing Heritability in sALS. Front. Neurosci. 14, (2020).

34. Improvements and impacts of GRCh38 human reference on high throughput sequencing data analysis. Genomics 109, 83–90 (2017).

35. Pan, B. et al. Similarities and differences between variants called with human reference genome HG19 or HG38. BMC Bioinformatics 20, (2019).

36. Miller, C. A. et al. Failure to detect mutations in U2AF1 due to changes in the GRCh38 reference sequence. (2021) doi:10.1101/2021.05.07.442430.

37. Li, H. et al. Exome variant discrepancies due to reference-genome differences. Am. J Hum. Genet. 108, 1239–1250 (2021).

38. Collins, R. L. et al. Author Correction: A structural variation reference for medical and population genetics. Nature 590, E55 (2021).

39. Quinlan, A. R. & Hall, I. M. BEDTools: a flexible suite of utilities for comparing genomic features. Bioinformatics 26, 841–842 (2010).

40. Li, H. Minimap2: pairwise alignment for nucleotide sequences. Bioinformatics 34, 3094–3100 (2018).

41. Krusche, P. et al. Best practices for benchmarking germline small-variant calls in human genomes. Nat. Biotechnol. 37, 555–560 (2019).

42. Van der Auwera, G. A. & O’Connor, B. D. Genomics in the Cloud: Using Docker, GATK, and WDL in Terra. (O’Reilly Media, 2020).

43. Farek, J. et al. xAtlas: Scalable small variant calling across heterogeneous next-generation sequencing experiments. doi:10.1101/295071.

44. Edge, P. & Bansal, V. Longshot enables accurate variant calling in diploid genomes from single-molecule long read sequencing. Nat. Commun. 10, 4660 (2019).

45. Shafin, K. et al. Haplotype-aware variant calling enables high accuracy in nanopore long-reads using deep neural networks. doi:10.1101/2021.03.04.433952.

46. Sahraeian, S. M. E. et al. Deep convolutional neural networks for accurate somatic mutation detection. Nat. Commun. 10, 1–10 (2019).

47. Walker, B. J. et al. Pilon: an integrated tool for comprehensive microbial variant detection and genome assembly improvement. PLoS One 9, e112963 (2014).

48. Martin, M. et al. WhatsHap: fast and accurate read-based phasing. doi:10.1101/085050.

49. Zook, J. M. et al. Extensive sequencing of seven human genomes to characterize benchmark reference materials. Scientific Data 3, 1–26 (2016).

50. Li, H. Aligning sequence reads, clone sequences and assembly contigs with BWA-MEM. arXiv [q-bio.GN] (2013).

51. Regier, A. A. et al. Functional equivalence of genome sequencing analysis pipelines enables harmonized variant calling across human genetics projects. Nat. Commun. 9, (2018).

52. Poplin, R. et al. Scaling accurate genetic variant discovery to tens of thousands of samples. doi:10.1101/201178.

53. Li, H. et al. The Sequence Alignment/Map format and SAMtools. Bioinformatics 25, 2078–2079 (2009).

54. Rausch, T. et al. DELLY: structural variant discovery by integrated paired-end and split-read analysis. Bioinformatics 28, i333–i339 (2012).

55. Cameron, D. L. et al. GRIDSS: sensitive and specific genomic rearrangement detection using positional de Bruijn graph assembly. Genome Res. 27, 2050–2060 (2017).

56. Layer, R. M., Chiang, C., Quinlan, A. R. & Hall, I. M. LUMPY: a probabilistic framework for structural variant discovery. Genome Biol. 15, R84 (2014).

57. Chen, X. et al. Manta: rapid detection of structural variants and indels for germline and cancer sequencing applications. Bioinformatics 32, 1220–1222 (2016).

58. Kronenberg, Z. N. et al. Wham: Identifying Structural Variants of Biological Consequence. PLoS Comput. Biol. 11, e1004572 (2015).

59. Jeffares, D. C. et al. Transient structural variations have strong effects on quantitative traits and reproductive isolation in fission yeast. Nat. Commun. 8, 14061 (2017).

60. De Coster, W., D’Hert, S., Schultz, D. T., Cruts, M. & Van Broeckhoven, C. NanoPack: visualizing and processing long-read sequencing data. Bioinformatics 34, 2666–2669 (2018).

61. Sedlazeck, F. J. et al. Accurate detection of complex structural variations using single-molecule sequencing. Nat. Methods 15, 461–468 (2018).

62. Jiang, T. et al. Long-read-based human genomic structural variation detection with cuteSV. Genome Biol. 21, 1–24 (2020).

63. Tarasov, A., Vilella, A. J., Cuppen, E., Nijman, I. J. & Prins, P. Sambamba: fast processing of NGS alignment formats. Bioinformatics 31, 2032–2034 (2015).

64. Faust, G. G. & Hall, I. M. SAMBLASTER: fast duplicate marking and structural variant read extraction. Bioinformatics 30, 2503–2505 (2014).

65. Poplin, R. et al. A universal SNP and small-indel variant caller using deep neural networks. Nat. Biotechnol. 36, 983 (2018).

