## Supplementary Figures, Notes, and Tables for "Towards a Comprehensive Variation Benchmark for Challenging Medically-Relevant Autosomal Genes"

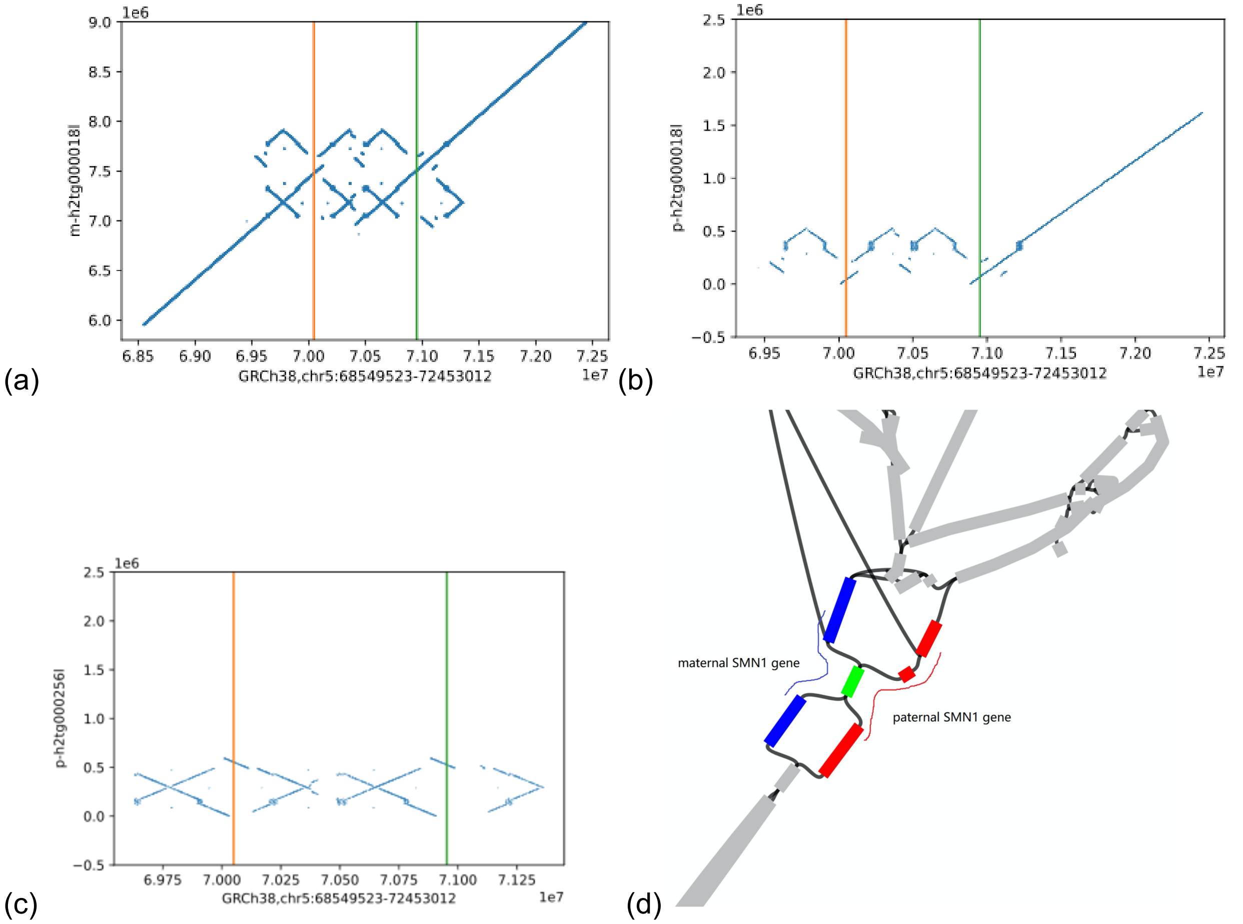


**Supplementary Figure 1:** Dot plots and assembly graph for HG002 assemblies in SMA region. (a) Maternal contig containing *SMN1* vs. GRCh38 in SMA region. (b) Paternal contig containing *SMN1* vs. GRCh38 in SMA region. (c) Paternal contig containing *SMN2* vs. GRCh38 in SMA region (*SMN2* was not assembled in the maternal contig). (d) Assembly graph for *SMN1* region, with maternal *SMN1* gene in blue and paternal *SMN1* gene in red.

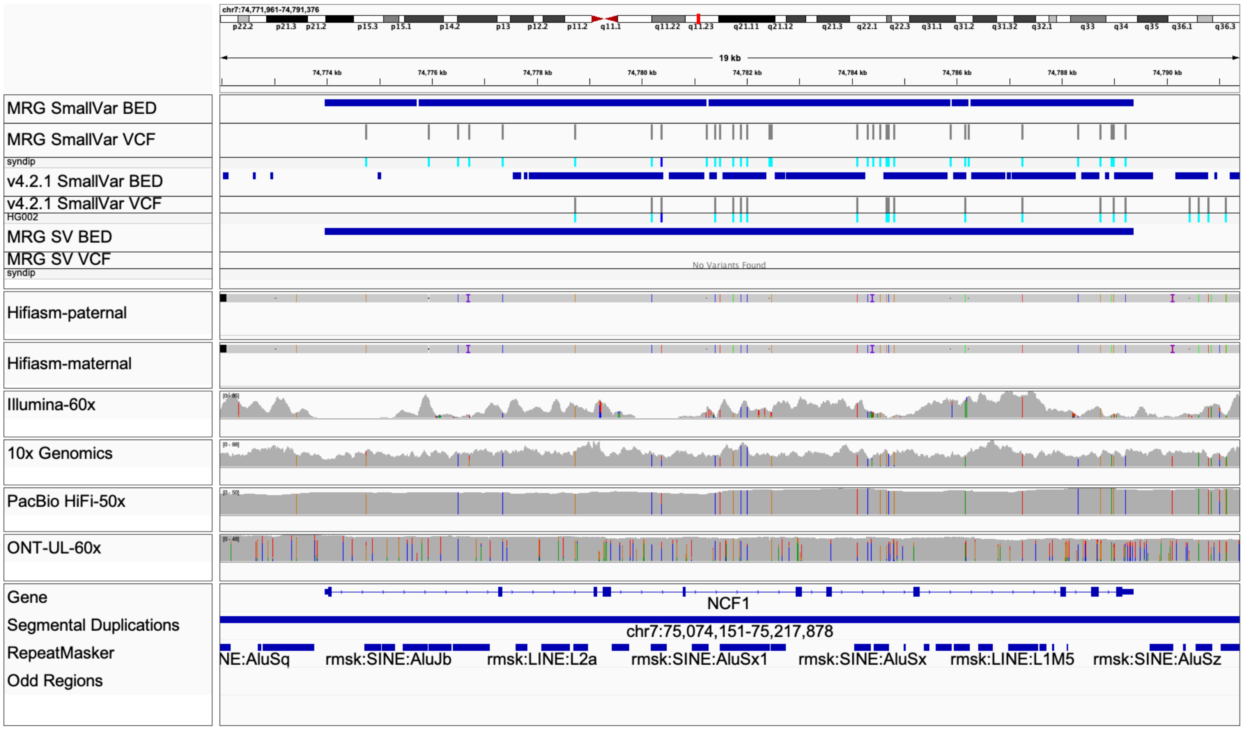


**Supplementary Figure 2:** Our new assembly-based benchmark almost completely resolves the gene *NCF1*. *NCF1* is in a 140 kb segmental duplication, resulting in some regions missing coverage by short reads, and the first 2 exons and some other variants missing from the v4.2.1 benchmark in GRCh38.

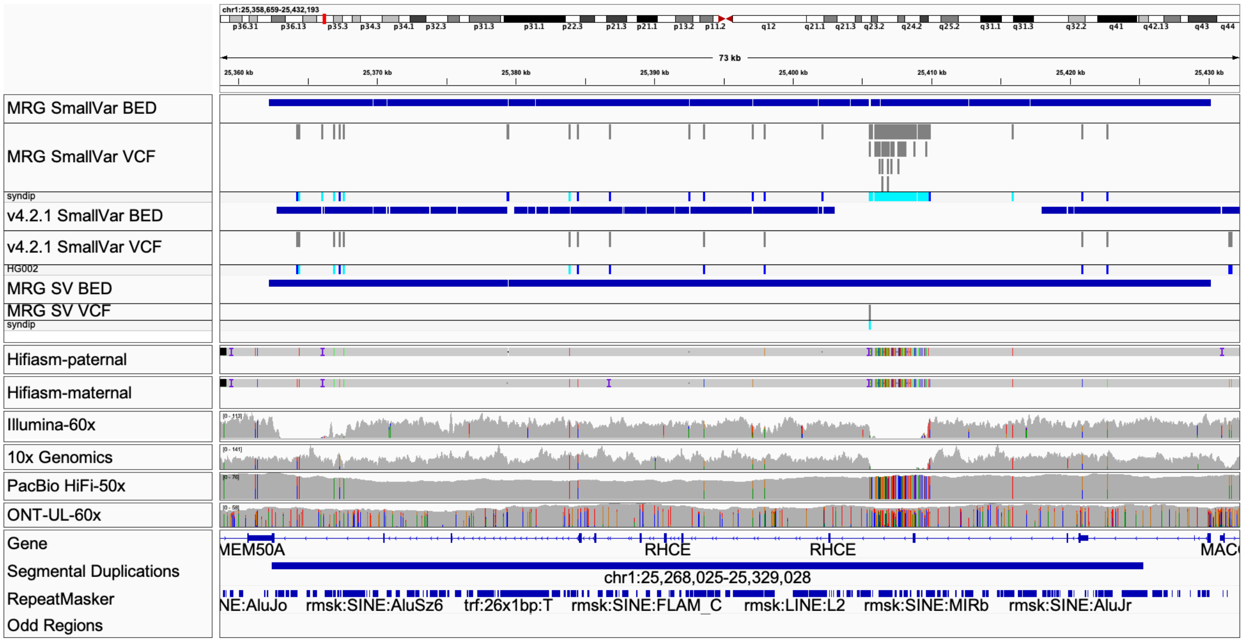


**Supplementary Figure 3:** The benchmark contains a 4.5 kb region of dense small variants and a structural variant in the gene *RHCE*, which may have resulted from a past gene conversion-like event with *RHD*. Short reads and linked reads from this region incorrectly map to *RHD* (GRCh38 shown).

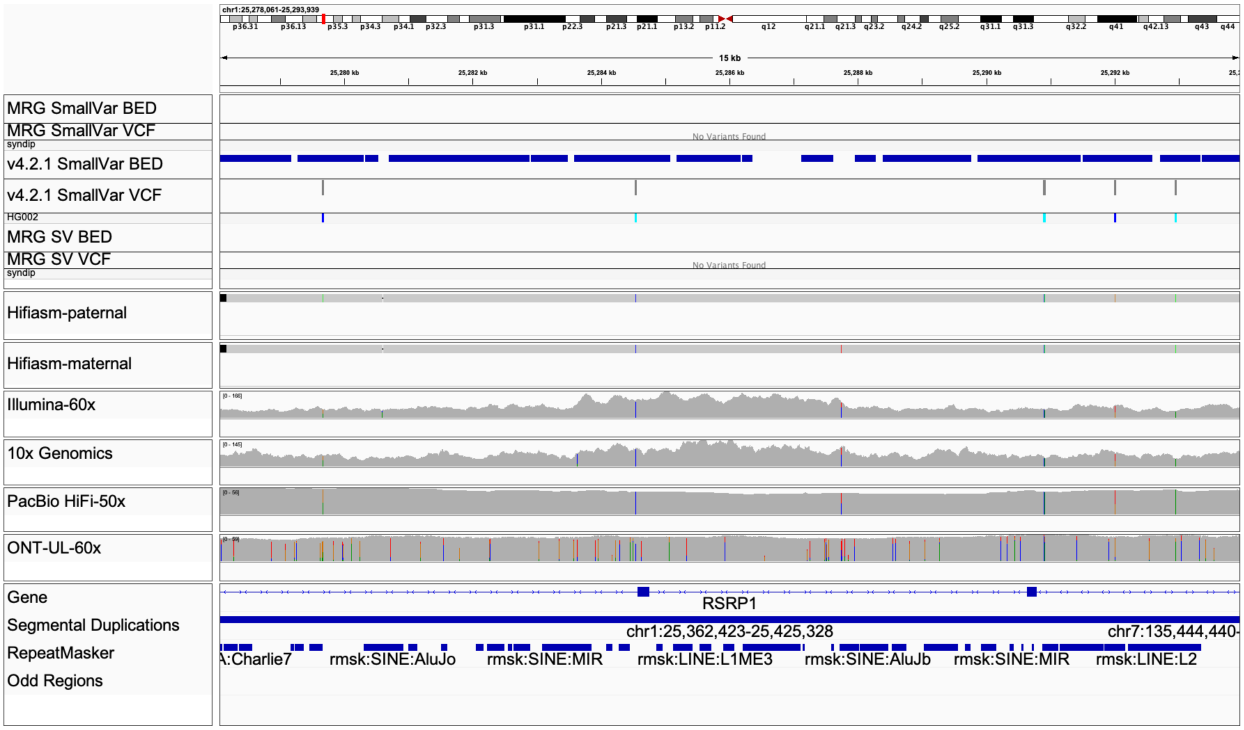


**Supplementary Figure 4:** An approximately 4.5 kb region in the segmentally duplicated gene *RHD* has about twice the normal coverage of short- and linked-reads due to reads mis-mapping from a gene conversion event in the homologous gene *RHCE* (GRCh38 shown). Note that the gene *RSRP1* overlaps with *RHD*, so it is in the label in the IGV session above.

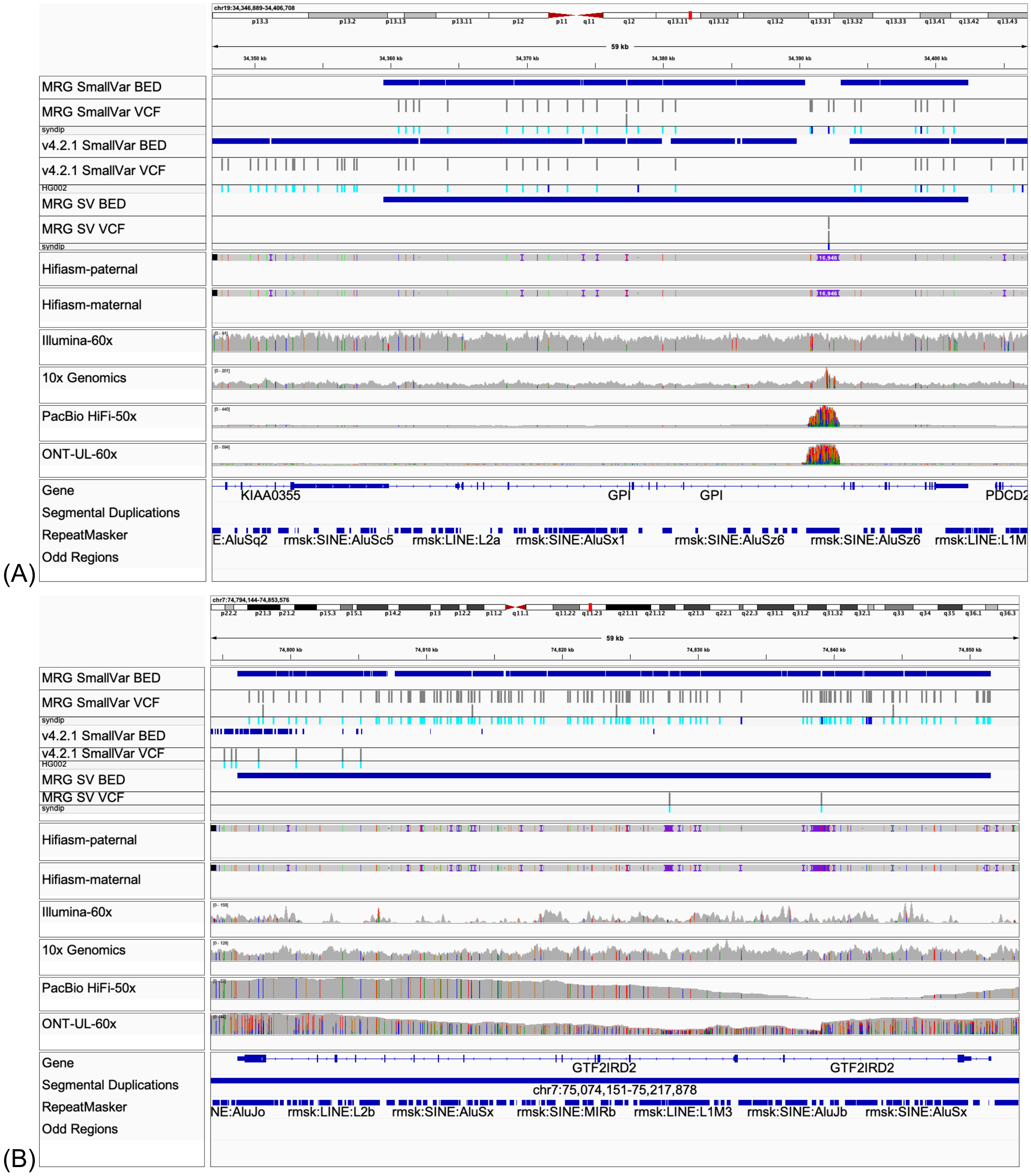


**Supplementary Figure 5: The new CMRG benchmark contains challenging SVs excluded from previous benchmarks.** (A) The new benchmark includes a 16,946 bp insertion in an intronic VNTR in the gene *GPI*. Even with long reads, this variant is challenging to call with current mapping-based SV callers. (B) The new benchmark includes two homozygous insertions in the segmentally duplicated gene *GTF2IRD2*. Many long reads containing the insertions align to the wrong copy of the segmental duplication, making it challenging to call SVs with mapping-based methods.

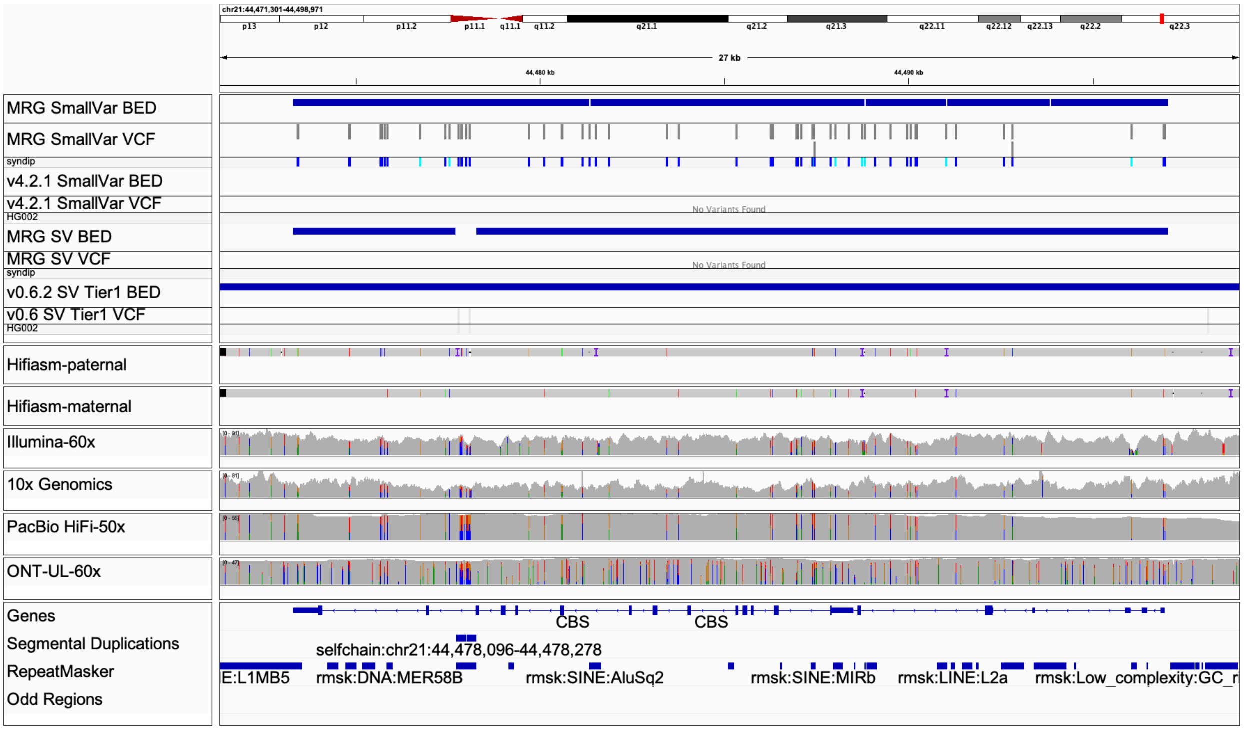


**Supplementary Figure 6**: Unlike GRCh38, variant calls across technologies are consistent with the benchmark on GRCh37 because GRCh37 does not contain the false duplication.

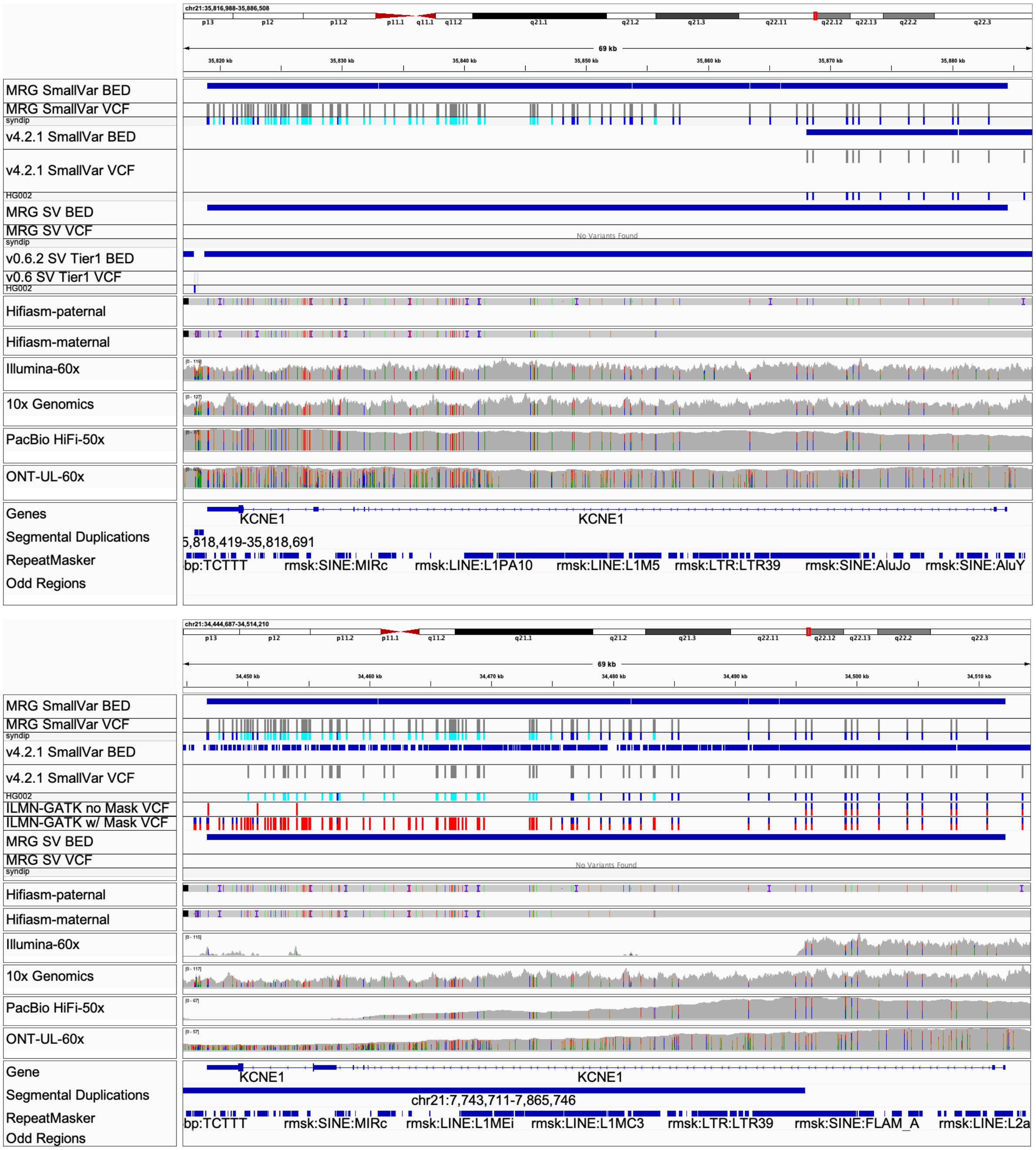


**Supplementary Figure 7: KCNE false duplication in GRCh38.** (A) Variant calls across technologies are consistent with the benchmark on GRCh37, because GRCh37 does not contain a false duplication. (B) GRCh38 contains a false duplication of part of the gene, so many reads mis-map to the false copy of the gene, *KCNE1B*

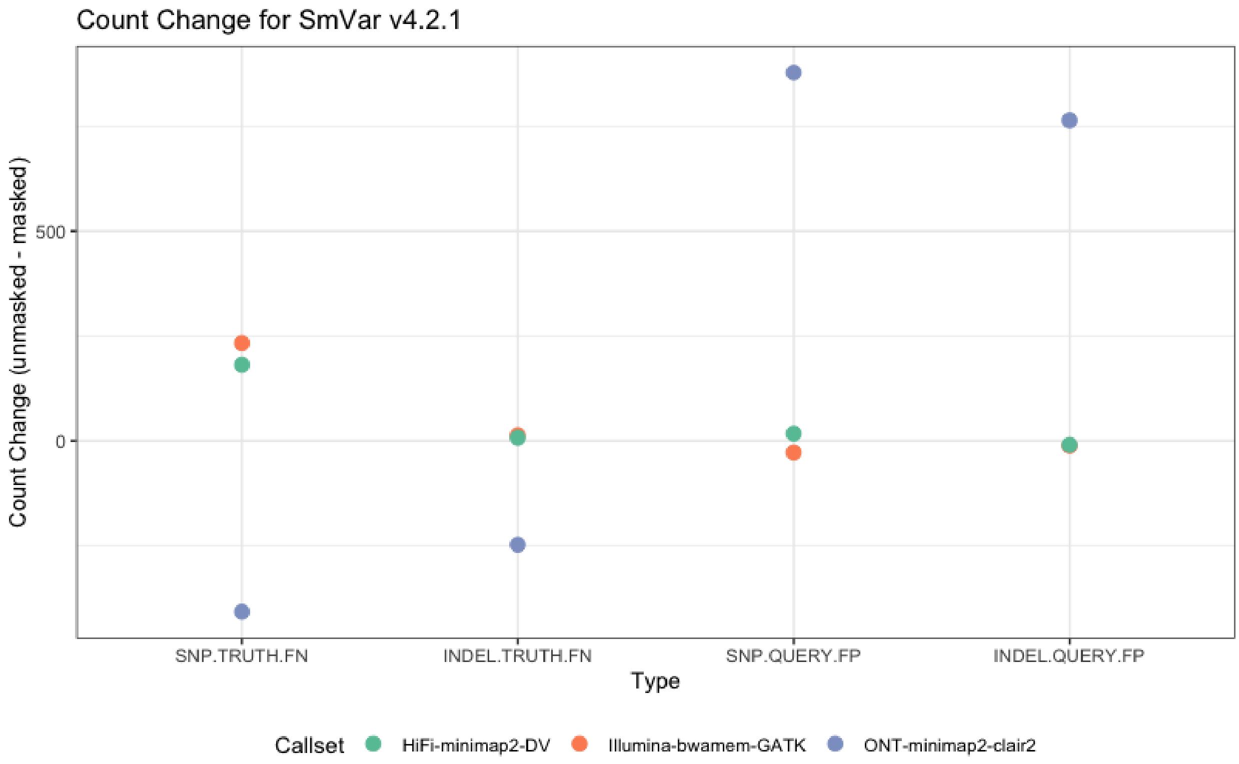


**Supplementary Figure 8:** Comparison of variant accuracy for GRCh38 before and after masking false duplications on chromosome 21. The new benchmark demonstrates only small changes to false positive and false negative counts for the whole genome benchmark v4.2.1 when mapping to the masked GRCh38, since v4.2.1 excludes most of the falsely duplicated regions.

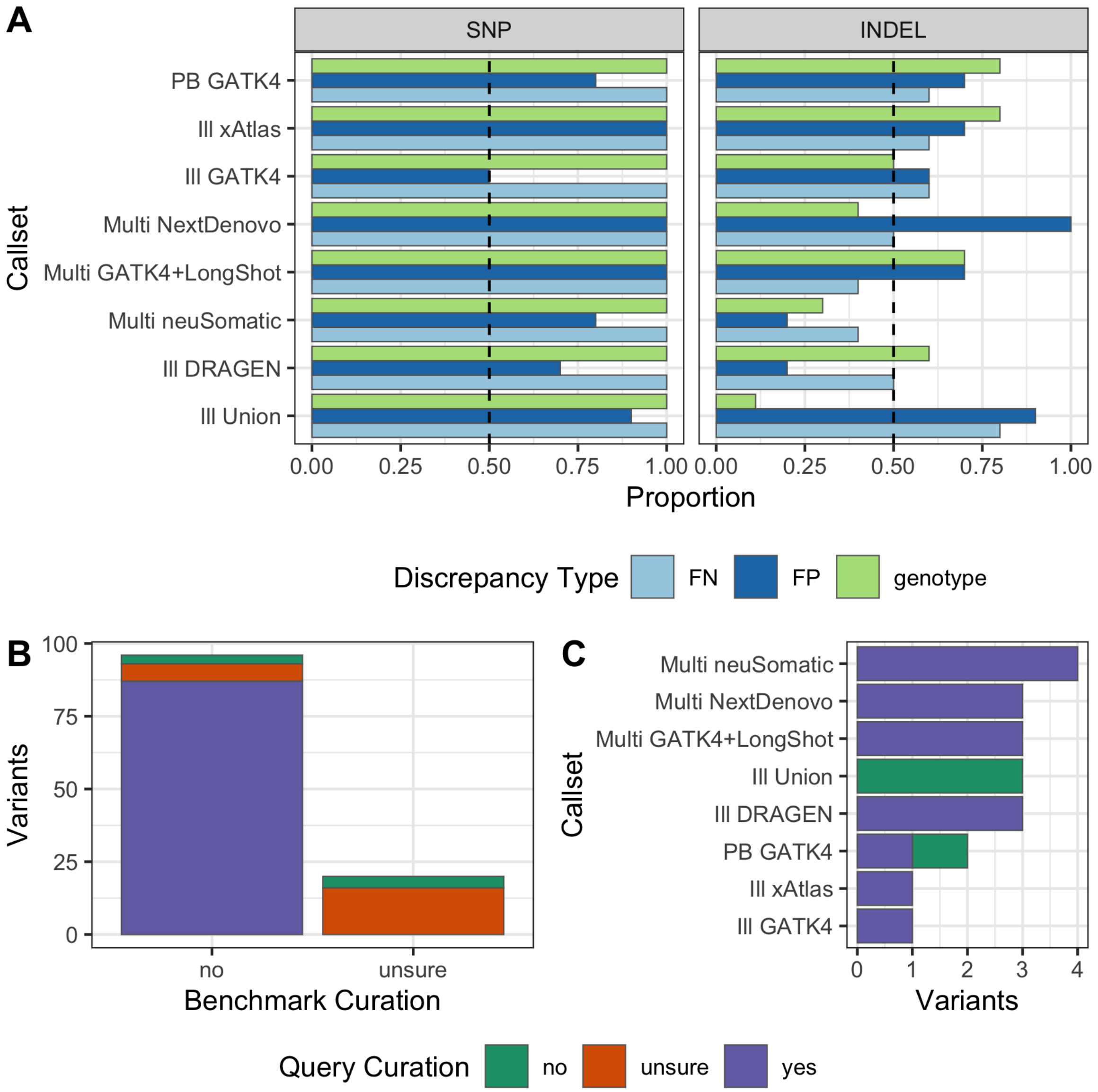


**Supplementary Figure 9: The v0.02.03 small variant CMRG benchmark reliably identified most types of errors, but some putative INDEL FPs and genotype errors needed to be excluded from v1.00.** We used hap.py with the vcfeval engine to compare each of the evaluation callsets against the v0.02.03 small variant CMRG benchmark. We later excluded all sites manually curated as unsure or error in the benchmark from the v1.00 small variant CMRG benchmark. A) The proportion of curated FP and FN variants by callset where the benchmark set was correct and the query callset was incorrect. The dashed black line indicates the majority threshold, 50%, because GIAB's goal for benchmarks is to exceed this threshold. Curated variants from both GRCh37 and GRCh38 (20 total) were used to calculate proportions. (B) Breakdown of the total number of variants by manual curation category, excluding variants from panel A where the benchmark was deemed correct and query incorrect, showing some sites were difficult to curate with current technologies. (C) Benchmark unsure variants by callset. 44/50 and 59/63 of the errors identified by the evaluation on GRCh37 and GRCh38, respectively, were excluded by curation of the common false positives and false negatives, so the accuracy of the v1.00 benchmark is higher. Technology abbreviations are: ONT=Oxford Nanopore, PB=PacBio HiFi, Ill=Illumina PCR-free, Multi= Combined ONT, PB, and Ill.

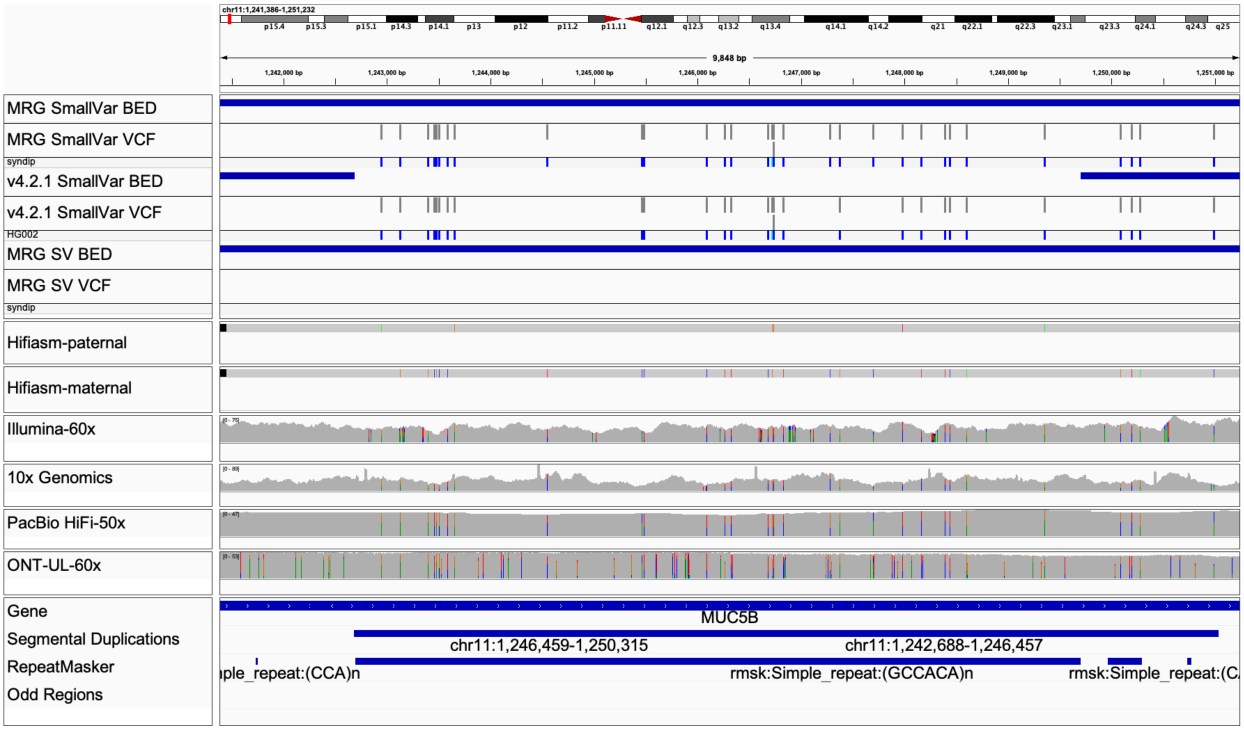


**Supplementary Figure 10:** The new benchmark contains a series of small variants in this long exonic tandem repeat in *MUC5B*. When there are many variants within a tandem repeat, they can often be called in a variety of ways. They may not be counted as true positives when benchmark if they are represented differently from the benchmark and partially called incorrectly or filtered. We created a new stratification to highlight false positives and false negatives in these regions in HG002.

(a)

(b)

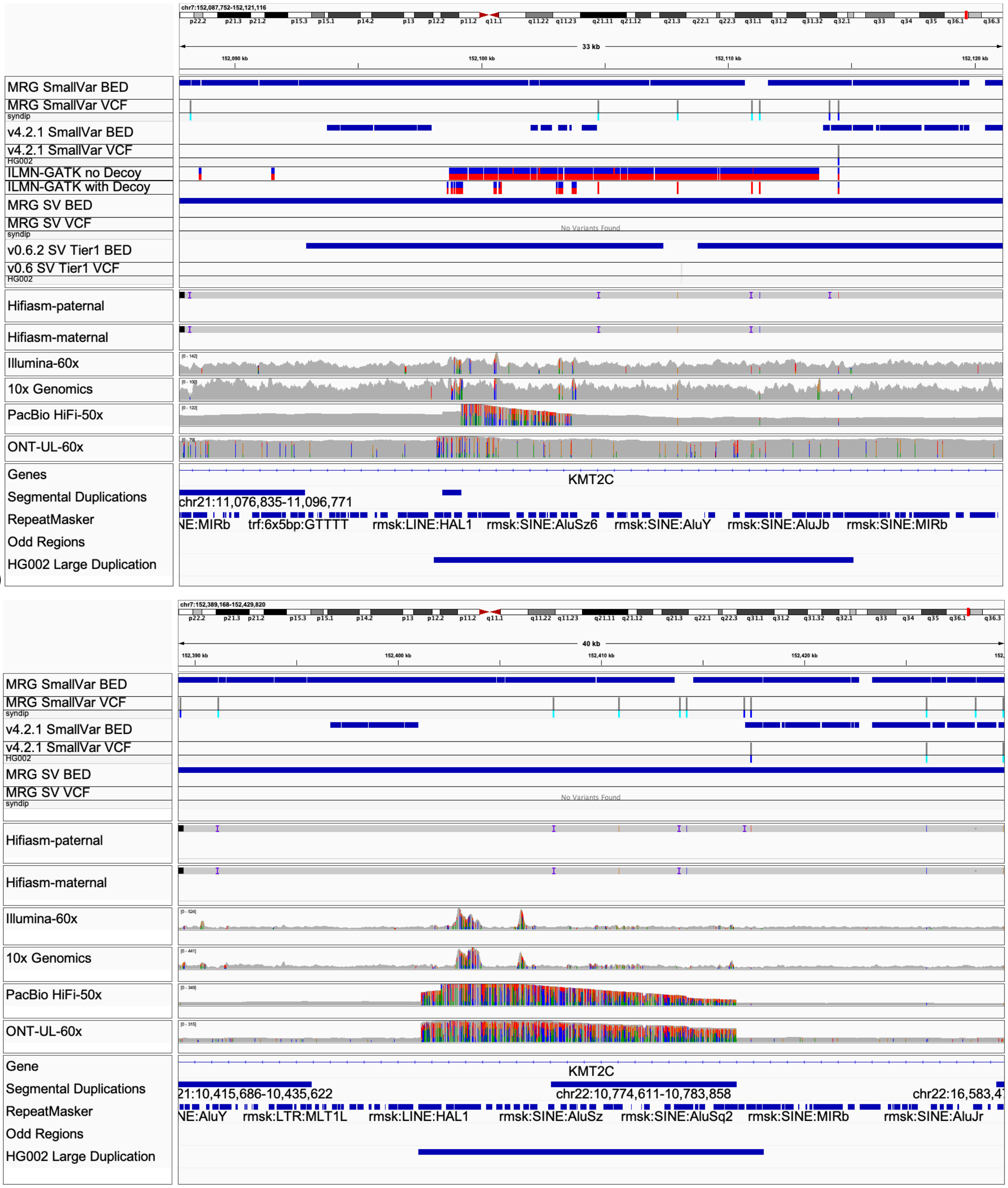


**Supplementary Figure 11**: We created a new stratification for a large, divergent duplication of part of the gene *KMT2C* in HG002 missing from GRCh38 (b) and only included in the hs37d5 decoy for GRCh37 (a). Mapping-based methods can call many false positives in this region in GRCh38 or in GRCh37 without the decoy, because reads from the duplication incorrectly map to the gene, as shown in the very high coverage and high density of variants in the alignments across technologies on GRCh38.

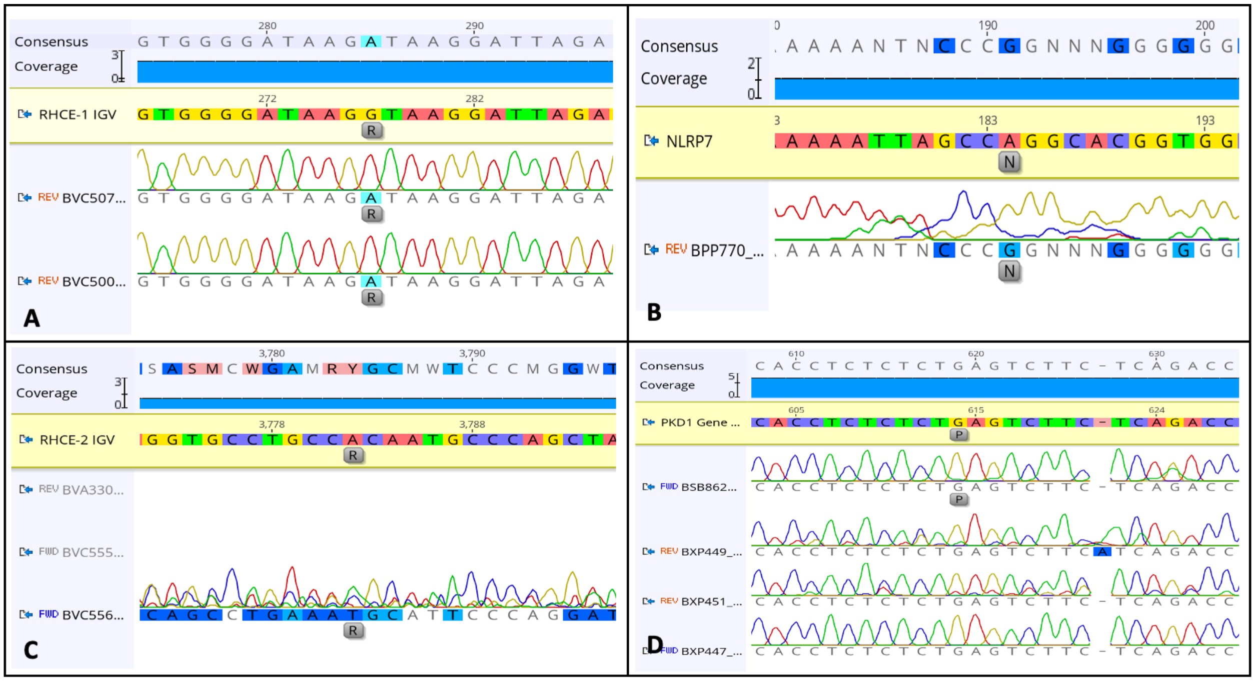


**Supplementary Figure 12: A**. *Variant Confirmed*. These variants have clean coverage by at least 1 primer, the base call is clearly variant at the expected base location, and all other base calls in the surrounding region clearly match the reference; **B**. *Variant Supported Not Confirmed*. These variants show support for a variant at the given base location, but other, unsupported variants are shown in the surrounding region as a result of noise in the chromatogram; **C**. *Variant Covered but not Confirmed*. These variants have coverage by at least 1 primer, but the sequence cannot be discerned by the Sanger trace. Usually this is a result of messy sequencing with multiple nucleotide peaks at all bases around the variant; **D**. *PKD1*. This variant does not show support for the G-->C variant identified by NGS. Several forward and reverse primers show a homozygous reference base call at this location.

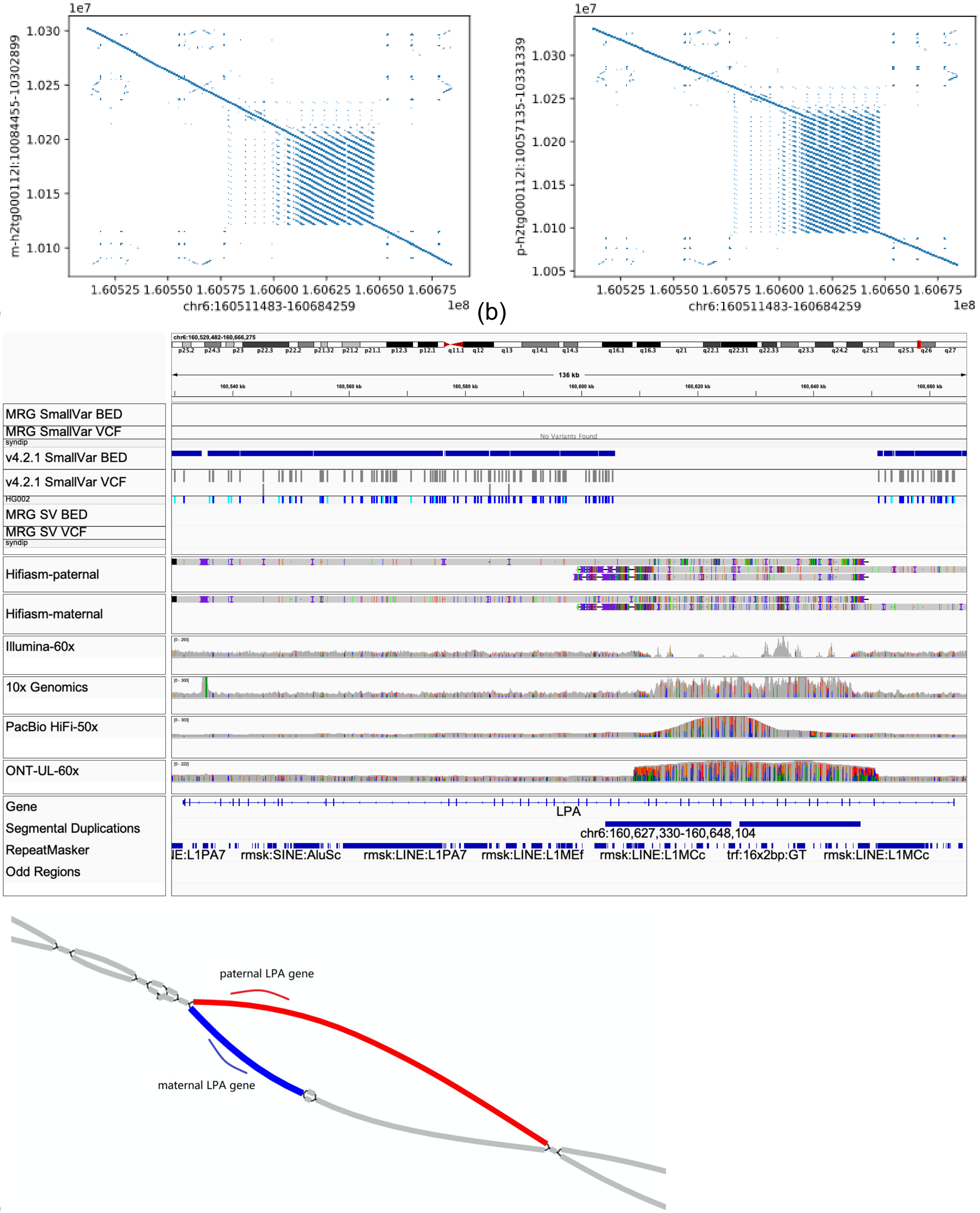


(a) (b) (c) (d)

**Supplementary Figure 13:** Dot plots, IGV, and assembly graph for HG002 assemblies in the medically important gene *LPA*, which contains 45 kb and 100 kb expansions of the tandemly duplicated kringle IV repeats relative to GRCh38. (a) Maternal contig containing 45 kb expansion vs. GRCh38. (b) Paternal contig containing 45 kb expansion vs. GRCh38. (c) IGV showing complex assembly and read alignments in repeat expansion. (d) Assembly graph for LPA region, with maternal LPA gene in blue and paternal LPA gene in red.

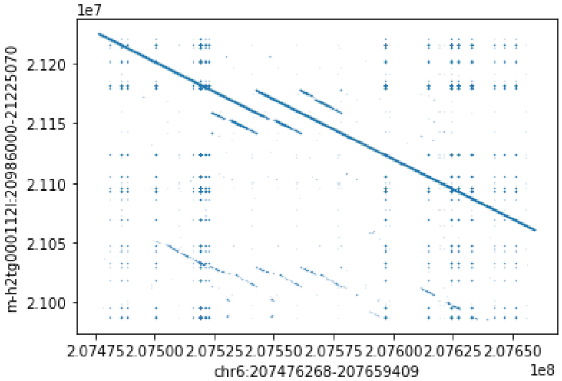

(a)

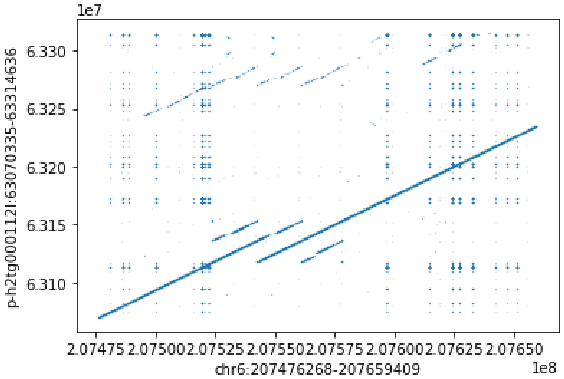

(b)

**Supplementary Figure 14:** Dot plots for HG002 assemblies in the medically important gene *CR1*, which contains 18 kb contractions of the large tandem repeats relative to GRCh38. (a) Maternal contig and (b) Paternal contig dot plots vs. GRCh38. These are excluded in the current benchmark due to breaks in the dipcall alignments.

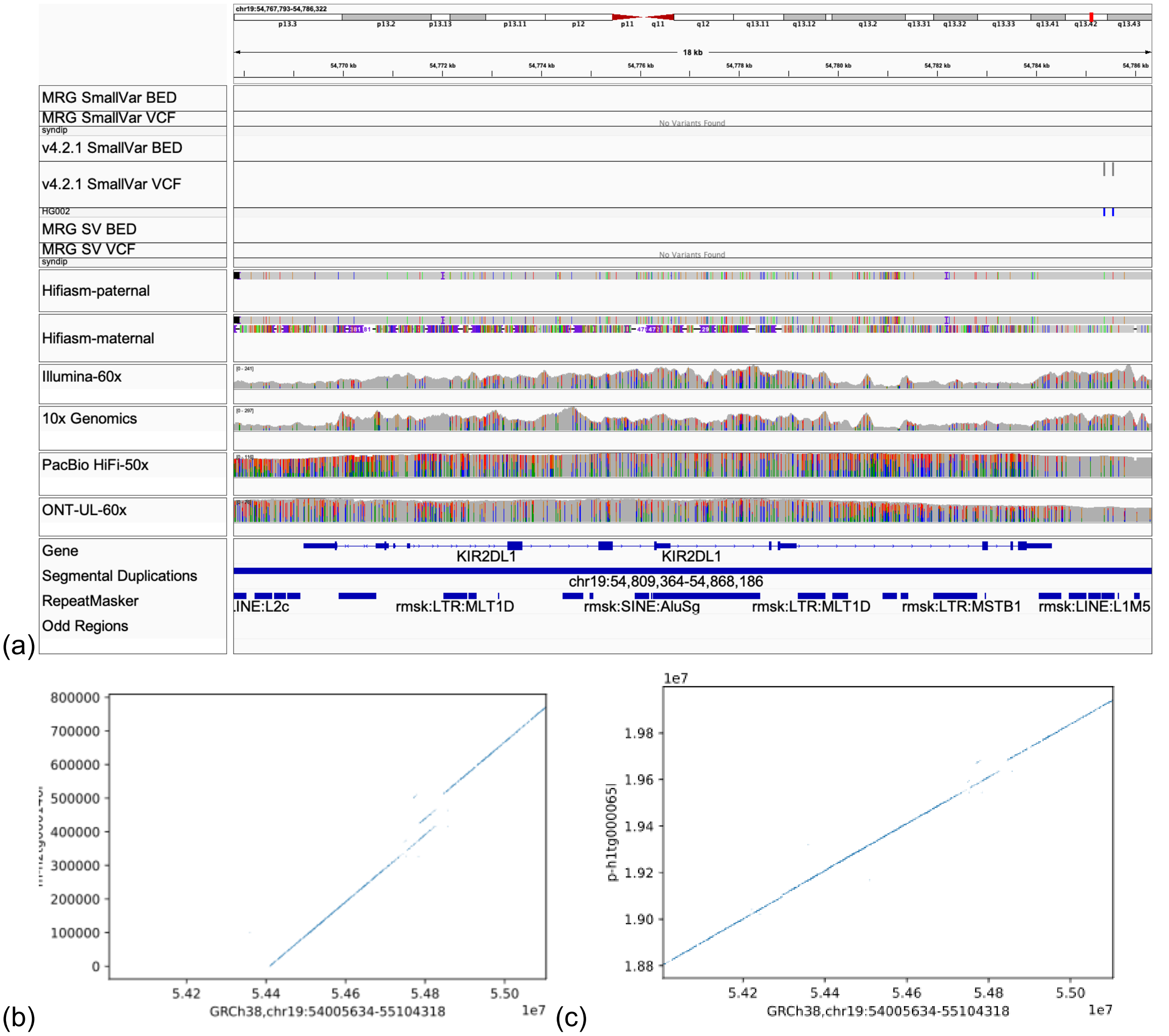


**Supplementary Figure 15:** (a) IGV screenshot showing high coverage of the gene *KIR2DL1.* (b) The high coverage results from the duplication of this gene on the maternal haplotype of HG002 relative to GRCh38 shown in the dot plot. (c) The paternal haplotype does not contain the duplication of this region.

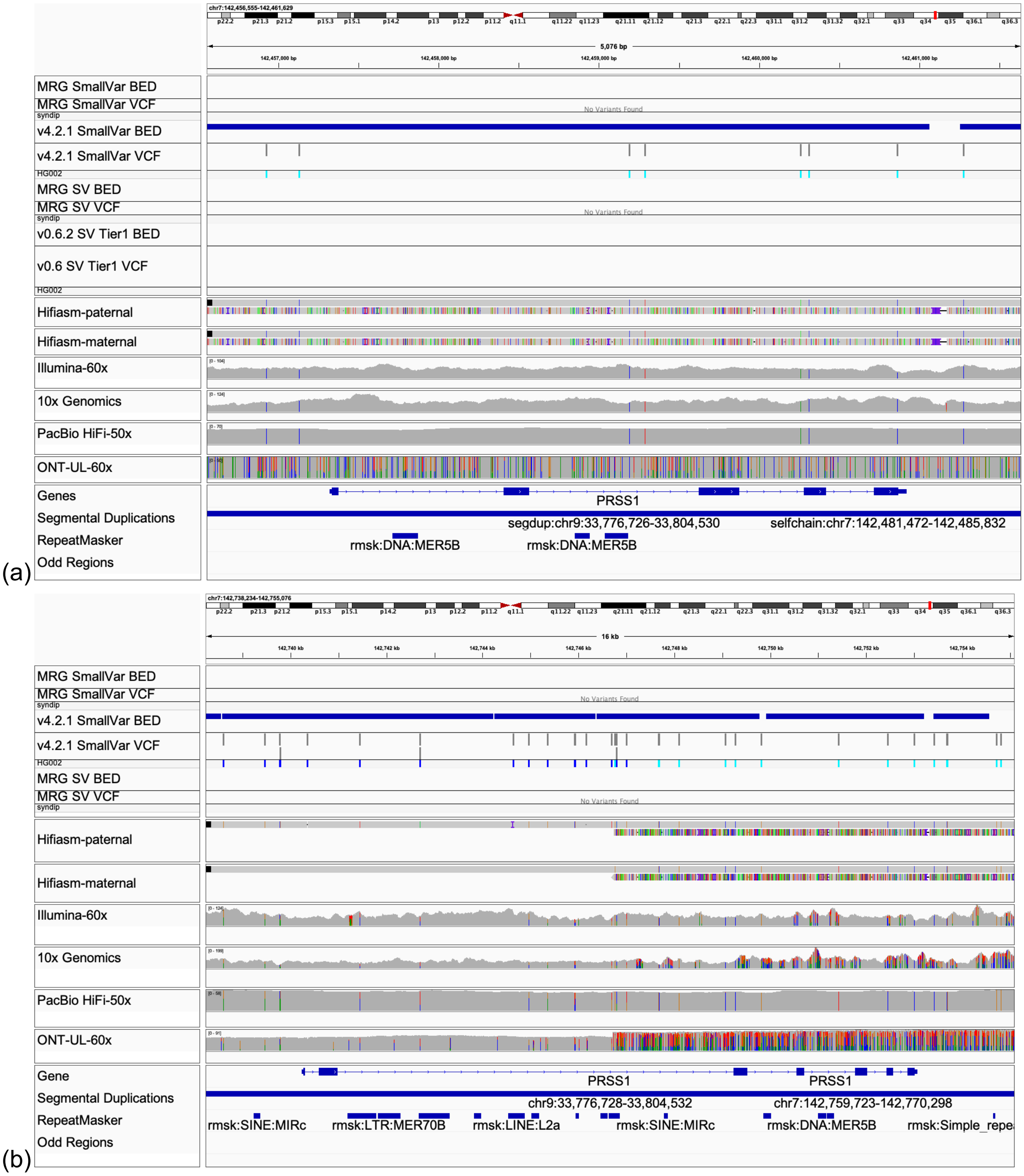


**Supplementary Figure 16**: The gene *PRSS1* has an extra copy in both haplotypes of HG002 relative to GRCh37 (a) and GRCh38 (b). GRCh37 has normal coverage because the extra copy is similar to the decoy sequence hs37d5, whereas GRCh38 only contains this sequence in the alternate locus NT_187562.1, which was not included in our reference for alignments.

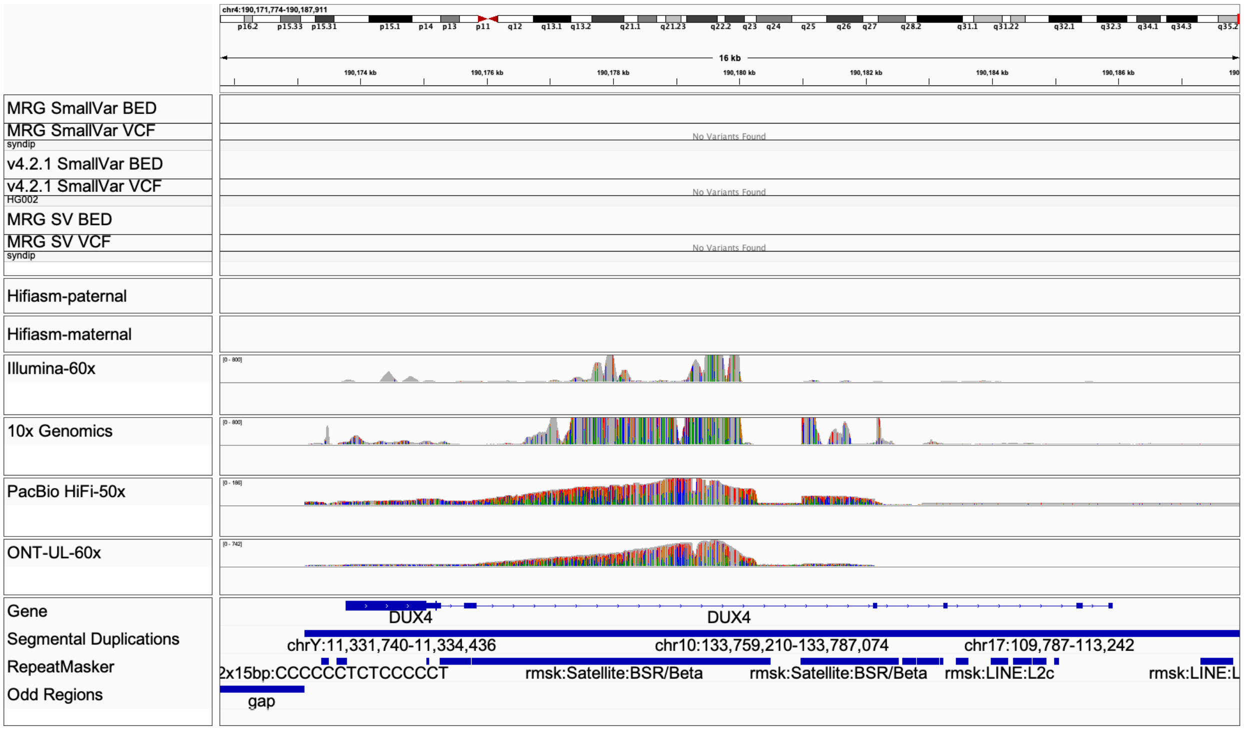


**Supplementary Figure 17:** Very high coverage of the gene *DUX4* due to many gaps and missing sequence in the D4Z4 region in GRCh38, missing extra copies of the gene that are in HG002. This region is not represented at all in GRCh37

### Supplementary Note 1: Manually excluded benchmark errors

Curation identified that the initial version of the benchmark contained some errors, which fell in a few categories: (1) regions of high homozygosity, where hifiasm sometimes misses one haplotype, mostly INDELs in homopolymers and dinucleotide tandem repeats (e.g., 1:6496542 on GRCh37/chr1:6436482 on GRCh38), (2) homopolymers and dinucleotide tandem repeats where the benchmark call is off by one or two bps, respectively, likely due to reduced HiFi read accuracy in long homopolymers particularly C/G homopolymers (e.g., 2:1148810 on GRCh37/chr2:1153124 on GRCh38; and 3:136053953 on GRCh37/chr3:136335111 on GRCh38). These are sometimes unclear during curation when PCR-free Illumina is also noisy or biased; (3) large insertions in homopolymers that result in homopolymers >20 bp on one or both haplotypes (e.g., 19:45419394 on GRCh37/chr19:44916137 on GRCh38); (4) a small number of hifiasm consensus errors that cause large INDELs or a series of small variants that are supported by a single read (e.g., 10:135235879 on GRCh37/chr10:133422375 on GRCh38); (5) a small number of adjacent insertion and deletion calls in the assembly alignment, for which a dipcall bug causes only the insertion to be called (e.g., 12:7355777/chr12:7203181 on GRCh37/GRCh38). When curating the homopolymers and dinucleotide tandem repeats, we trusted the PCR-free Illumina reads that traversed the entire repeat region if there was no evidence of mapping errors (e.g., in some segmental duplications) or systematic sequencing errors (e.g., at some G/C homopolymers). In v1.00, we remove all 215 regions with benchmark errors or unclear calls identified by manual curation on GRCh37 or GRCh38.

We evaluated small complex variants in tandem repeats longer than 100 bp in a different way. These variants can be represented in VCF in many ways (e.g. as one line or multiple lines), are often partially filtered by mapping-based methods, and can be very challenging to curate in a genome browser due to different representations in aligned reads from different mappers/technologies. Because assembly-based methods usually perform best in these regions, we compared variants from a trio-hicanu assembly to our hifiasm-based benchmark small variants, because they use different consensus approaches. Variant calls and genotypes agreed between the two methods for 1,343/1,361 SNVs and 944/1,004 INDELs in these complex tandem repeats. Upon curation of the 21 variants unique to the trio-hicanu assembly, we found that most were errors in the benchmark, so we excluded all complex repeats that had any variants unique to the trio-hicanu assembly. Upon curation of the 53 remaining differences on GRCh38, variants with different genotypes or variants unique to the benchmark, we found that 20 were correct in the benchmark (often due to hicanu missing a haplotype), 17 were unclear (often due to C/G homopolymers), and 16 were errors in the benchmark (mostly due to noise in the HiFi reads in homopolymers and dinucleotide tandem repeats).

We evaluated the SV benchmark by comparing four callsets: (1) pbsv from PacBio HiFi alignments, (2) assembly-based calls using ONT and Illumina, (3) a union of 5 Illumina-based callers, and (4) a union of 2 ONT-based callers. We found the benchmark reliably identified false positives and false negatives across all 4 callsets. Upon manual curation only two sites were identified as problematic: a 50 bp net insertion that was represented as two smaller insertions in the tandem repeat in the benchmark, and a 376 bp deletion in the benchmark that was represented in many ONT alignments as two smaller deletions in the tandem repeat. Although these are correct in the benchmark, current SV benchmarking tools fail to compare these different representations, so we exclude these regions from the v1.0 SV benchmark bed. In addition, we found that some callers represent tandem duplications and tandem repeat expansions as duplications with SVTYPE=DUP (and sometimes incorrectly as translocations or inversions), whereas our benchmark calls these as insertions with the annotation REPTYPE=DUP. These can be counted as true positives by ignoring SVTYPE (--type-ignore -p 0 option in truvari) or by changing SVTYPE from DUP to INS in the query VCF. We also compared Bionano optical mapping-based SV calls to the 50 benchmark SVs >=500 in size. Because many of these SVs were near the limit of detection of optical mapping, we curated these calls, and all were supported by the Bionano data.

### Supplementary Note 2: New Stratifications

To enable users to better understand performance in new types of challenging regions included in this CMRG benchmark, we created new stratifications for regions where some variant callers may have particularly high error rates relative to the benchmark. The first new stratification is a 16 kb region in an intron of the gene *KMT2C* that has multiple copies in HG002 that are not present in GRCh38 (GRCh37 has similar problems in *KMT2C*, but to a lesser degree when using the hs37d5 decoy sequence). Mis-mapped reads, particularly from long reads, can cause a cluster of many false positives using mapping-based approaches, causing 277 of the 386 false positives in the HiFi-DeepVariant callset, as shown in **Supplementary Figure 11.** Similarly, additional small regions in *KMT2C* have high coverage by short reads, causing false positives. The second new stratification is for complex variants in tandem repeats longer than 100 bp. When more than one variant occurs in a long tandem repeat, the complex variant can often be represented in many different ways. Mapping-based methods sometimes filter part of the complex variant, which can cause the variants in the tandem repeat to be counted as many false positives and false negatives. These contribute to a significant fraction of all false positives and false negatives for most methods (e.g., 28% of errors for HiFi-DeepVariant and 24% of errors for Illumina-GATK). **Supplementary Figure 10** shows an example of a complex variant in a tandem repeat in a coding region of *MUC5B*.

### Supplementary Note 3: Medically Relevant Genes not yet benchmarked

The hifiasm assembly resolved both haplotypes of *LPA*, which is an important gene often related to cardiovascular diseases, but it is excluded in the CMRG benchmark due to very large insertions that cause a break in contig alignments (**Supplementary Figure 13)**. The general structure of *LPA* contains multiple tandemly duplicated copies of the same region (ie. kringle IV repeats of ~8 kb). These repeats often range between 10 and 50 repeats that are transcribed and translated[28,29]. The overall repeat number was associated with cardiovascular disease risk and is thus important to resolve correctly. The HG002 hifiasm assembly resolved the entire *LPA* region, including the kringle IV repeats, which have a total length of 7.2 kb in GRCh38. We confirmed that the 44.1 kb and 99.9 kb insertion sizes from hifiasm for the maternal and paternal haplotypes, respectively, were consistent with the insertions predicted by an independent trio-phased Bionano optical mapping assembly (45.0 kb and 101.2 kb). This complex, large expansion of the kringle IV repeats can be represented in many different ways in a VCF with different levels of precision (e.g., as a large insertion, a tandem duplication, or a CNV, and the copies may differ or include small variants). Existing benchmarking tools cannot compare these different representations robustly, partly limited by the VCF format[30]. To benchmark assemblies of this gene in HG002, the sequences could be compared directly to the hifiasm contigs, which we have annotated for *LPA* and other genes using LiftOff[31]. *CR1*, a gene implicated in Alzheimer's disease[8], is similarly resolved by hifiasm but contains a large SV that causes a break in the dipcall/minimap2 alignment (**Supplementary Figure 14)**. For *CR1* in the GRCh38 region chr1:207538089-207573740, the reference allele is 35.6 kb in length. Both paternal and maternal alleles of HG002 are 17.1 kb in length, or a 18.5 kb homozygous deletion, which is consistent with the Bionano deletion prediction of 18.6 kb for both alleles.

Other genes are excluded from the benchmark because they have extra copies in HG002 but not in GRCh38. For example, genes in the KIR region are highly variable and CNVs are observed frequently in the population, with 35 alternate loci and 15 novel patches in GRCh38.p13. Hifiasm resolves the paternal allele in a single contig, but the maternal allele is split into 3 contigs in the KIR region. The maternal allele has an extra copy of the gene *KIR2DL1* that is tandemly duplicated, so that minimap2 aligns both copies to the region, and this duplication is supported by alignments of short-, linked-, and long-reads (**Supplementary Figure 15**). There is no standard way to represent or benchmark small variants in duplicated regions, so we excluded these in our benchmarks. In addition, further work will be needed to fully resolve the maternal haplotype in a single contig in the KIR region. Similarly, the medically important gene *PRSS1* has an extra divergent copy in both HG002 alleles that is similar to an alternate locus in GRCh38 and to the decoy sequence hs37d5 for GRCh37 (**Supplementary Figure 16**). *DUX4* in the D4Z4 locus is also excluded in the CMRG benchmark because it is not well-represented in GRCh38 or GRCh37, with many gaps, and the gene has very high coverage due to multiple copies of the gene in HG002 but not in GRCh38 (**Supplementary Figure 17**).

**Supplementary File 1 High Priority Clinical Gene lists.xlsx:** Additional information about high priority clinical genes.

**Supplementary File 2 MRG_stats_table_GRCh38.tsv:** Overlaps of the 5,038 genes on GRCh38 primary assembly between both HG002 GRCh38 v4.2.1 and HG002 hifiasm v0.11.

**Supplementary File 3 HG002_v0.11_benchmark-v4.2.1.summary.csv**: Benchmarking of the hifiasm v0.11 assembly-based variants called with dipcall against the GIAB v4.2.1 benchmark for HG002

**Supplementary File 4 HG002 CMRG small variant evaluations.xlsx**: Tables summarizing all benchmarking statistics against CMRG benchmark and evaluation callsets

**Supplementary File 5 Manual curation results for evaluation and common errors in v0.02.03 small variant benchmark**

**Supplementary File 6 LongRangePCR_Sanger_Confirmation_Procedures:** Primer designs and reaction conditions for Long-Range PCR and Sanger Confirmation.

**Supplementary File 7 remaining genes excluded from CMRG.tsv**: Genes excluded from the CMRG benchmarks, with likely reasons for exclusion annotated for GRCh38 in the last column

**Supplementary File 8 pipeline.38_nodecoy.txt**: Commands for BWA-GATK variant calling on normal GRCh38 reference

**Supplementary File 9 pipeline.38_nodecoy_mask.txt**: Commands for BWA-GATK variant calling on v1 masked GRCh38 reference

**Supplementary Table 1:** Results of Long Range PCR and Sanger Sequencing to confirm variants in genes with segmental duplications in the new HG002 medically relevant gene benchmark.

| Gene | Long Range PCR Amplicon Size (bp) | Total Variants | Number of Variants Definitively Confirmed | Number of Variants With No Primer Coverage | Number of Variants Supported (Not Definitively Confirmed) | Number of Variants Covered but Noisy Sanger traces | Number of Variants Contradicted | Reasons for variants not definitively confirmed |
|---|---|---|---|---|---|---|---|---|
| *CBS* | 12,216 | 28 | 19 | 5 | 3 | 0 | 0 | 3 variants fall in a long tandem repeat with a complex variant; the 1 insertion and 1 deletion in the benchmark are likely equivalent to the Sanger and HiFi representation of a series of SNVs |
| *DCLRE1C* | 12,543 | 8 | 5 | 3 | 0 | 0 | 0 | N/A |
| *FCGR2B* | 14,703 | 38 | 34 | 2 | 2 | 0 | 0 | N/A |
| *FLG* | 7,625 | 27 | 24 | 3 | 0 | 0 | 0 | N/A |
| *FXN* | 3,984 | 29 | 23 | 0 | 5 | 1 | 0 | Messy sequencing doesn't allow for call of variant |
| *GTF2IRD2* | 21,791 | 51 | 24 | 21 | 2 | 4 | 0 | Messy sequencing doesn't allow for call of variant |
| *HYDIN* | 3,392 | 1 | 0 | 0 | 1 | 0 | 0 | Long homopolymer shows variant at location but sequencing around it is messy |
| *NCF1* | | 4 | 4 | 0 | 0 | 0 | 0 | N/A |

| | | | | | | | | |
|---|---|---|---|---|---|---|---|---|
| | 6,466 | | | | | | | |
| *NLRP7* | 12,067 | 54 | 24 | 15 | 6 | 9 | 0 | Messy sequencing doesn't allow for call of variant |
| *PKD1* exons 2-7 | | 1 | 1 | 0 | 0 | 0 | 0 | N/A |
| *PKD1* exons 8-12 | | 2 | 1 | 0 | 0 | 0 | 1 | 1 variant from benchmark is not supported by either of the 2 primers |
| *PKD1* exons 22-26 | | 2 | 2 | 0 | 0 | 0 | 0 | N/A |
| PKD1 exons 27-34 | | 5 | 5 | 0 | 0 | 0 | 0 | N/A |
| *RHCE* region 1 | 6,226 | 85 | 41 | 32 | 7 | 5 | 0 | Messy sequencing doesn't allow for call of variant |
| *RHCE* region 2 | 10,000 | 18 | 18 | 0 | 0 | 0 | 0 | N/A |
